# The influence of gene flow on population viability in an isolated urban caracal population

**DOI:** 10.1101/2023.07.20.549918

**Authors:** Christopher C. Kyriazis, Laurel E.K. Serieys, Jacqueline M. Bishop, Marine Drouilly, Storme Viljoen, Robert K. Wayne, Kirk E. Lohmueller

**Author notes:** deceased 26 December 2022 (Robert K. Wayne). This paper is dedicated to the memory of Robert Wayne for his innumerable contributions to mammalian conservation and evolutionary genetics.

## Abstract

Wildlife populations are becoming increasingly fragmented by anthropogenic development. Such small and isolated populations often face an elevated risk of extinction, in part due to inbreeding depression. Here, we examine the genomic consequences of urbanization in a caracal (*Caracal caracal*) population that has become isolated in the Cape Peninsula region of the city of Cape Town, South Africa and is thought to number ∼50 individuals. We document low levels of migration into the population over the past ∼75 years, with an estimated rate of 1.3 effective migrants per generation. As a consequence of this isolation and small population size, levels of inbreeding are elevated in the contemporary Cape Peninsula population (mean F_ROH>1Mb_=0.20). Inbreeding primarily manifests as long runs of homozygosity >10Mb, consistent with the effects of isolation due to the rapid recent growth of Cape Town. To explore how reduced migration and elevated inbreeding may impact future population dynamics, we parameterized an eco-evolutionary simulation model. We find that if migration rates do not change in the future, the population is expected to decline only slightly, with a low projected risk of extinction. However, if migration rates decline or anthropogenic mortality rates increase, the potential risk of extinction is greatly elevated. To avert a population decline, we suggest that translocating migrants into the Cape Peninsula to initiate a genetic rescue may be warranted in the near future. Our analysis highlights the utility of genomic datasets coupled with computational simulation models for investigating the influence of gene flow on population viability.

## Introduction

Anthropogenic development has resulted in pervasive habitat destruction and fragmentation (Haddad et al., 2015; Wilson et al., 2016). Consequently, formerly large and widespread populations are increasingly being transformed into smaller, fragmented populations. These small and isolated populations face an elevated risk of extinction due to a multitude of factors, among which is the deterioration of fitness due to increased genetic drift and inbreeding (Caughley, 1994; Frankham, 2005). In particular, inbreeding depression, or the loss of fitness due to inbreeding, may pose a key threat to population viability in small populations (Benson et al., 2016; Hedrick & Garcia-Dorado, 2016; Johnson et al., 2010; Keller & Waller, 2002; Robinson et al., 2019). Compared to the more gradual degradation of fitness resulting from a fixation of weakly deleterious alleles due to increased genetic drift in small populations (i.e., mutational meltdown; (Lynch, Conery, & Burger, 1995)), inbreeding depression driven by recessive strongly deleterious mutations can drive relatively rapid fitness declines in cases where inbreeding is rapid and severe (Kyriazis, Wayne, & Lohmueller, 2021; Robinson, Kyriazis, Yuan, & Lohmueller, 2023). Thus, minimizing close inbreeding in natural populations by maintaining connectivity and ample population sizes is vital for averting genetic threats to extinction.

For small populations that are derived from historically-large populations, the risk of local extinction due to inbreeding depression may be especially high (Angeloni, Ouborg, & Leimu, 2011; Hedrick & Garcia-Dorado, 2016; Kyriazis et al., 2021; Mathur & DeWoody, 2021; Oosterhout, 2019; Robinson et al., 2022, 2023, 2019; Van Der Valk, Manuel, Marquez-Bonet, & Guschanski, 2019). Recessive strongly deleterious mutations, the primary driver of inbreeding depression, can accumulate in large and genetically diverse populations, where they tend to be hidden as heterozygotes from negative selection (Hedrick, 2002; Hedrick & Garcia-Dorado, 2016; Kyriazis et al., 2021; Robinson et al., 2019; Szpiech et al., 2019; Van Der Valk et al., 2019). Following a decline to small population size (*N_e_*<50), inevitable close inbreeding can expose recessive strongly deleterious mutations as homozygotes, potentially resulting in severe inbreeding depression and further population decline. An illustrative example of these dynamics comes from mountain lions, a species with a large historical effective population size on the order of *N_e_*=100,000 that has been greatly contracted and fragmented due to anthropogenic development and hunting, resulting in a growing number of small and isolated populations (Saremi et al., 2019). Most famously, the highly isolated Florida panther population nearly declined to extinction in the 1990s due to severe inbreeding depression (Roelke, Martenson, & O’Brien, 1993). Extinction was averted only after migrant individuals from Texas were introduced, resulting in a ‘genetic rescue’ that included a dramatic increase in population size from ∼20 to >100 individuals (Johnson et al., 2010; van de Kerk, Onorato, Hostetler, Bolker, & Oli, 2019).

In the wake of the success of this and several other genetic rescues, there has been a growing call to more widely implement assisted gene flow as a management tool for small and isolated populations (Bell et al., 2019; Fitzpatrick, Mittan-Moreau, Miller, & Judson, 2023; Ralls et al., 2018; Whiteley, Fitzpatrick, Funk, & Tallmon, 2015). Nevertheless, genetic rescue initiatives remain rare, with only ∼20 documented instances to date (Fitzpatrick et al., 2023; Frankham, 2015). One potential limitation of this management strategy is that the beneficial effects of genetic rescue in the form of heterosis (i.e., increased fitness in hybrids due to masking of recessive deleterious alleles; (Whitlock, Ingvarsson, & Hatfield, 2000)) are likely to be short-lived, particularly when the recipient population is unable to grow (Bell et al., 2019; Kyriazis et al., 2021; Lotsander et al., 2021; Pérez-Pereira, Caballero, & García-Dorado, 2022; Robinson et al., 2023). This is because inevitable inbreeding in small populations will expose introduced recessive deleterious mutations as homozygotes soon after genetic rescue has occurred, leading to renewed inbreeding depression. However, recent theoretical work has suggested that the long-term effectiveness of genetic rescue can potentially be improved by a careful consideration of the demographic history and load of recessive deleterious variation in the source population (Kyriazis et al., 2021; Pérez-Pereira et al., 2022). Specifically, these studies suggest that targeting moderate-sized source populations that are somewhat purged of recessive deleterious variation can result in a lowered threat of inbreeding depression and extinction in the recipient population under future inbreeding (Kyriazis et al., 2021; Pérez-Pereira et al., 2022). Thus, obtaining a detailed understanding of the demographic history of the source and recipient population may be an essential component of maximizing the effectiveness of genetic rescue.

In this study, we employ a genomic resequencing dataset and computational simulation modelling to examine the relationship between gene flow and population viability in an urban caracal (*Caracal caracal*) population in the city of Cape Town, South Africa. Caracals are elusive medium-sized felids with a large geographic range that encompasses much of Africa, extending into the Middle East and South Asia, though are most abundant in South Africa (Avenant et al., 2016; Veals, Burnett, Morandini, Drouilly, & Koprowski, 2020). Caracals are present the Cape Peninsula of Cape Town – the site of Table Mountain National Park, a UNESCO World Heritage Site recognized for its rich biodiversity – where they have become isolated by the rapid growth of the city over the past several decades (Fig. 1; Rebelo, Holmes, Dorse, & Wood, 2011; Turok & Borel-Saladin, 2014). By one population density estimate (Avenant et al., 2016), there may be approximately 48 caracals in the Cape Peninsula (Leighton et al., 2022). Moreover, GPS data have shown that very little migration between the peninsula and nearby populations is occurring (Fig. S1; Leighton et al., 2022; Serieys et al., 2023). Thus, the large historical population size of caracals combined with the recent isolation of the small Cape Peninsula population suggests that it may be at an especially high risk of extinction due to inbreeding depression. Moreover, Cape Town’s caracals also face numerous additional threats to persistence including incidental poisoning from rodenticides, exposure to persistent organic pollutants and metals, vehicle collisions, poaching, disease, and lethal management (Leighton et al., 2022; Parker et al., 2023; Serieys et al., 2019; Viljoen et al., 2020). Thus, a pressing question for the Cape Peninsula caracal population is whether it will be able to persist without interventions to reduce anthropogenic mortality rates and bolster migration, or if it may soon be threatened by population decline and local extinction.

**Figure 1:**
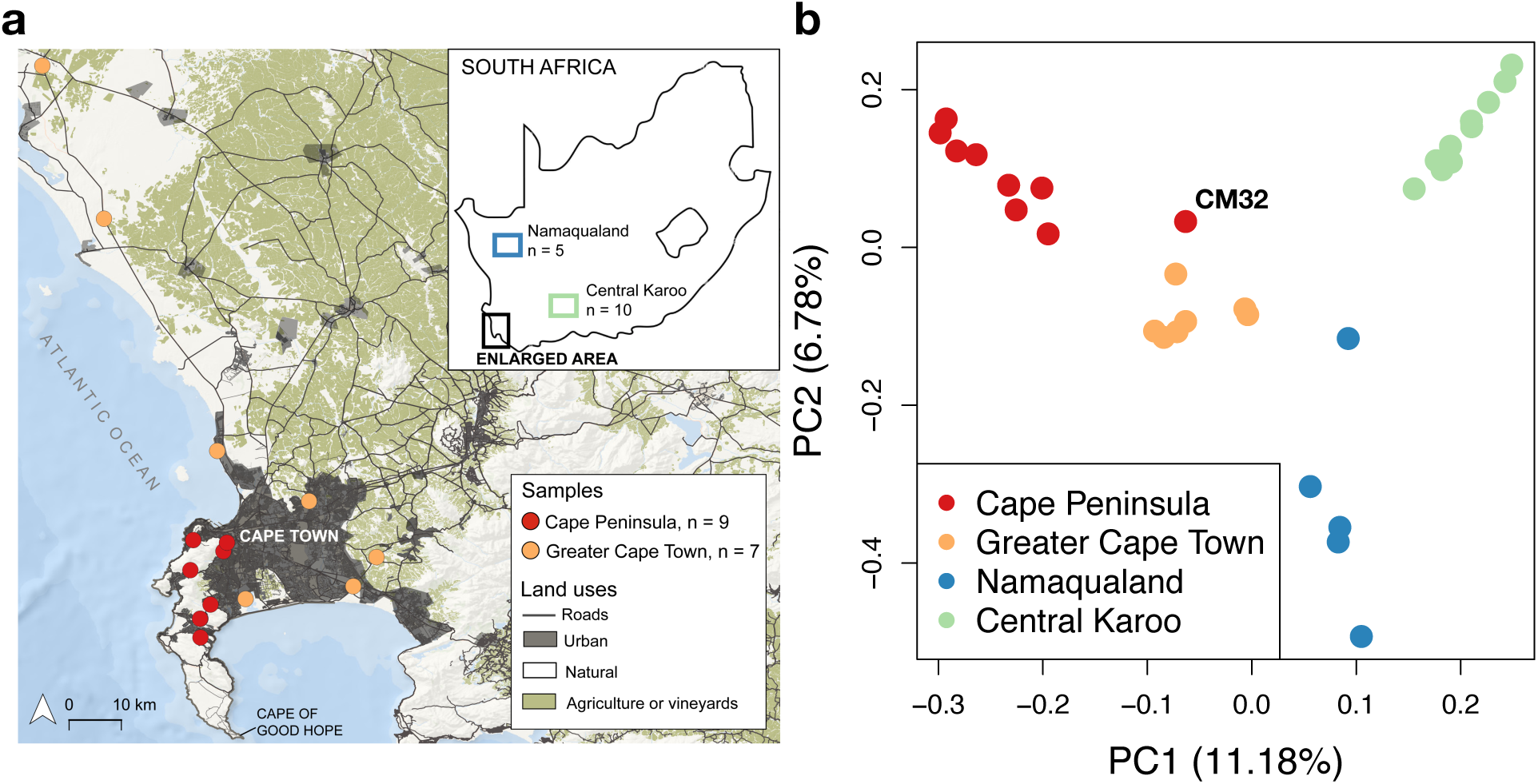
Map of sampling localities and characterization of population structure using principal component analysis. (a) Map of the Cape Peninsula and the Greater Cape Town area showing localities of sampled individuals, including an inset map of South Africa showing Namaqualand and Central Karoo regions. See Figure S1 for a more detailed map of Cape Town including GPS collar locations for 25 caracals. (b) Principal component analysis of all 31 sampled individuals color-coded by source population. Note that each population separates out with the exception of CM32, an individual sampled in the Cape Peninsula that clustered with the Greater Cape Town population.

## Methods

### Sampling & NGS sequencing

We sampled caracals from four populations: the Cape Peninsula (CP; n=9), the Greater Cape Town region (GCT; n=7), the Central Karoo (CK; n=10), and Namaqualand (NMQ; n=5; Fig. 1). We selected these individuals to focus sampling on the Cape Peninsula and Greater Cape Town populations, while also including non-urban populations that are thought to be large and are less fragmented (Avenant et al., 2016; Veals et al., 2020) as a point of comparison. Caracals in CP were primarily sampled within Table Mountain National Park (TMNP), which encompasses approximately 320 km^2^ of fragmented, natural habitat isolated by approximately 800 km^2^ of City of Cape Town (CoCT) urban matrix. The GCT includes the CoCT urban matrix and surrounding natural areas. The GCT is thus a mosaic of urban, light industrial, agricultural, and natural reserves both within and on the outskirts of CoCT (Fig. 1; CoCT mean human population density: 1530 people/km^2^). The CK is a rural region located approximately 350 km north-east of GCT, whereas NMQ is a rural region located approximately 480 km north of GCT. Both CK and NMQ are semi-desert regions that consist mainly of privately-owned extensive small-stock farmland with exceptionally low human population densities (CK: 1.8 people/km^2^; NMQ: 1.1 people/km^2^). Croplands are rare and restricted to small areas immediately adjacent to farm houses whereas stock largely comprises free-ranging sheep that feed on native vegetation. For individuals sampled in the CP or the GCT, blood and skin tissue was collected from caracals that were captured using cage traps (detailed in Serieys et al., 2023), or opportunistically recovered mortalities. In the CK and NMQ, muscle or skin tissue was collected from dead caracals killed by hunters or farmers during standard culling operations (CK: Cape Nature sampling permits 0056-AAA007-00161 and AAA007-00074-0056; NMQ: no permit required to sample postmortem individuals). Animal handling in the CP and GCT was approved by the University of Cape Town Animal Ethics Committee (2014/V20/LS), Cape Nature (AAA007-0147-0056), and SANParks (2014/CRC/2014-017, 2015/CRC/2014-017, 2016/CRC/2014-017, 2017/CRC/2014-017).

DNA was extracted from samples using Qiagen Blood and Tissue kits. Library preparation and sequencing of 150bp paired-end reads was completed at the Vincent J. Coates Genomics Sequencing Laboratory at University of California, Berkeley and MedGenome using an Illumina NovaSeq (Table S1).

### Data processing

To process raw reads, we adapted a pipeline from the Genome Analysis Toolkit (GATK) (Van der Auwera et al., 2013) Best Practices Guide. We used BWA-MEM (Li, 2013) to align paired-end 150bp raw sequence reads to the domestic cat genome (felCat9.0)(Buckley et al., 2020), followed by removal of low-quality reads and PCR duplicates. In the absence of a caracal reference genome, we opted to use this high-quality reference genome given the high degree of synteny that is observed in felids (Armstrong et al., 2022; Samaha, Wade, Mazrier, Grueber, & Haase, 2021). As we do not have a database of known variants, we did not carry out Base Quality Score Recalibration, but instead carried out hard filtering of genotypes (see below).

We used GATK HaplotypeCaller to perform joint genotype calling at all sites, including invariant sites. We filtered genotypes to include only high-quality biallelic SNPs and monomorphic sites, removing sites with Phred score below 30 and depth exceeding the 99^th^ percentile of total depth across samples. In addition, we removed sites that failed slightly modified GATK hard filtering recommendations (QD < 4.0 || FS > 12.0 || MQ < 40.0 || MQRankSum < −12.5 || ReadPosRankSum < −8.0 || SOR > 3.5), as well as those with >25% of genotypes missing or >75% of genotypes heterozygous. We masked repetitive regions using a mask file downloaded from https://www.ncbi.nlm.nih.gov/assembly/GCF_000181335.3/, generated using RepeatMasker (Smit, Hubley, & Green, n.d.). Finally, we applied a per-individual excess depth filter, removing genotypes exceeding the 99^th^ percentile of depth from a distribution calculated for each individual, as well as sites having fewer than six reads for each individual.

### Population structure

To characterize population structure among sampled caracal populations, we employed several different approaches implemented within SNPrelate v1.14 (Zheng et al., 2012). These included performing principal component analysis (PCA), constructing a tree using hierarchical clustering based on identity-by-state (IBS), and estimating kinship coefficients for all samples. For these analyses, we pruned SNPs for linkage disequilibrium (ld.threshold=0.2) and filtered out sites with minor allele frequency below 0.05, resulting in 49,941 SNPs for analysis. Finally, we also used SNPrelate to estimate genome-wide F_ST_ among populations. We employed the Weir & Cockerham approach for estimating F_ST_ (Weir & Cockerham, 1984).

### Genetic diversity & inbreeding

We calculated heterozygosity for each individual by computing the proportion of called sites with heterozygous genotypes. To visualize patterns of genetic diversity across individual genomes, we calculated heterozygosity in non-overlapping 1 Mb windows in the autosomal portion of each sampled genome. We removed windows in the bottom 5% tail of the number of calls per window, as such windows often have high variance in heterozygosity.

To assess the extent of inbreeding across samples, we infererd runs of homozygosity (ROH) in autosomal regions of the genome using BCFtools/RoH (Narasimhan et al., 2016). We used the -G30 flag and allowed BCFtools to estimate allele frequencies. We calculated individual inbreeding coefficients (F_ROH_) by summing the total length of all ROH calls >1 Mb and dividing by 2329.7 Mb, the autosomal genome length for the domestic cat reference genome.

To estimate the age of each ROH (i.e., time to coalescence) and determine at what time in the past inbreeding occurred, we employed a model where the length of ROH decays exponentially via recombination through time (Browning, 2008; Thompson, 2013). In this model, the number of generations to coalescence for an ROH (*g*) is given as *g*=100/(2\**L*), where *L* is the length of an ROH in centimorgans (cM). Following Saremi et al. (2019), we assumed an average recombination rate from the domestic cat of 1.1 cM per Mb as estimated by (Dumont & Payseur, 2008).

### Inferring ancestral and contemporary effective population sizes and migration rates

We used ∂a∂i (Gutenkunst, Hernandez, Williamson, & Bustamante, 2009) to infer demographic models from the site frequency spectrum (SFS) of neutral variants in our genomic data set. Specifically, our aim was to estimate the parameters of a divergence model for the Cape Peninsula and Greater Cape Town populations including historical and contemporary effective population sizes and migration rates. Given our somewhat limited sample sizes, we focused on fitting a relatively simple two-epoch divergence model with symmetric migration.

We began by generating joint two-dimensional neutral SFS from the Cape Peninsula and Greater Cape Town populations. To generate a set of putatively neutral variants, we extracted variants from genomic regions that were >10kb from coding regions that did not overlap with repetitive regions. Additionally, we excluded un-annotated highly conserved regions that are under strong evolutionary constraint, identified by aligning the remaining regions against the zebra fish genome using BLASTv2.7.1 (Camacho et al., 2009) and removing any region which had a hit above a 1e-10 threshold. We used EasySFS (https://github.com/isaacovercast/easySFS) to then generate a folded neutral SFS for these regions using a hypergeometric projection to account for missing genotypes. We found that the number of SNPs in each SFS was maximized when projecting to 2n=12 for the Cape Peninsula population and 2n=10 for the Greater Cape Town population. These SFS were then used as input data for inferring demographic models in ∂a∂i.

A key challenge in estimating demographic models with recent population size changes, as might be expected for these caracal populations, is disentangling the timing and magnitude of population size decline. For instance, patterns in the SFS indicative of population decline can often be explained by a model with a very recent and severe decline, or more ancient and modest decline (Beichman et al., 2023; Keinan, Mullikin, Patterson, & Reich, 2007). Given this challenge, we chose to fix the duration of the most recent epoch in all of our demographic models to *t_div_*=25 generations, thus assuming that population divergence began in 1940 (i.e., 75 years before sampling in 2015, assuming a generation time of 3 years; see below). This assumption was informed by two sources of information: (1) rapid pace of human population growth in Cape Town and elsewhere in South Africa starting in the 1940s (Rebelo et al., 2011), a main factor driving habitat fragmentation and potentially population decline, and (2) evidence for abundant medium and long ROH >1Mb in the Cape Peninsula and Greater Cape Town populations, with a mean ROH age of ∼32.5 years in the Cape Peninsula (see Results), suggestive of recent population declines.

With this assumption of *t_div_*=25 generations, we proceeded to fit a divergence model for the Cape Peninsula and Greater Cape Town populations. This model assumes a constant size for the ancestral population, followed by split with symmetric migration. Thus, we estimate parameters of *N_anc_*, *N_CP_*, *N_GCT_*, and *m.* For this inference, we assumed a mutation rate of 8.6e-9 mutations per site per generation, informed by a recent estimate for domestic cats (Wang et al., 2022), and, in the absence of a generation time estimate for caracals, assumed a generation time of 3 years, as informed by our simulation model (see below). To perform inference, we permuted the starting parameter values and conducted 50 runs for each model. For each run, we calculated the log-likelihood and selected the maximum likelihood estimate. Finally, we used the mutation rate of 8.6e-9 mutations per site per generation and total sequence length (*L*) to calculate the diploid ancestral effective population size as *N_anc_*=ϴ/(4\**μ*\**L*). We scaled other inferred population size parameters by *N_anc_* and time parameters by 2\**N_anc_* to obtain values in units of diploids and numbers of generations.

### Overview of eco-evolutionary simulation model

To explore the potential impact of isolation and small population size on population viability in the Cape Peninsula population, we parameterized eco-evolutionary simulations using the non-Wright-Fisher (nonWF) models in SLiM v4.0.1 (Haller & Messer, 2016, 2019, 2023). The nonWF model allows for simulating genome-scale genetic variation together with ecological population dynamics, enabling an assessment of how an accumulation of deleterious variation may influence the fitness and viability of the simulated population.

In the nonWF model, each individual has an absolute fitness ranging from 0 to 1 that is determined by the selective and dominance effects of deleterious mutations in their genome. Fitness is assumed to be multiplicative across sites, thus assuming no epistasis. To model density dependence in the simulated population, the nonWF model employs a carrying capacity (*K*), which is used to rescale fitness during each year by the ratio of *K*/*N* (i.e., if *N*>*K* after reproduction, absolute fitness will be scaled down during subsequent viability selection). Note that population sizes in this model are therefore census population sizes, in contrast to the typical convention in population genetics of modelling effective population sizes. To model varying survival rates for individuals of different age, the nonWF model also allows for enforcing age-specific fitness rescaling. The resulting rescaled absolute fitness determines an individual’s probability of annual survival, as well as probability of reproduction (see below). See Figure S2 for a schematic of events that occur during each year of the simulation model.

### Life history parameters

Each year the nonWF model begins with an opportunity for mature males and females to reproduce (Fig. S2). Based on available information on caracals, we assume that females reach reproductive maturity at age 1 and males at age 2 (Veals et al., 2020). We further assume that every adult female has an 80% chance of reproducing each year, multiplied by their unscaled absolute fitness. For instance, an adult female with absolute fitness of 0.9 would have a 0.8*0.9=0.72 probability of initiating reproduction each year. Reproduction occurs for each female by randomly selecting an adult male as a partner. We assume the litter size for each mating event to be Poisson distributed with a mean of 3, capped at a maximum of 6, as available evidence suggests that caracal litter sizes are typically between 2-4 kittens, though sometimes up to 6 (Veals et al., 2020). Note that ∼5% of such Poisson draws will result in a litter size of 0, further reducing the overall probability of successful reproduction.

At the conclusion of each year, viability selection occurs, where individuals survive with a probability given by their rescaled absolute fitness. As noted above, fitness is rescaled based on individual age to model varying survival rates for different classes of individuals. We assumed age-specific fitness rescaling factors of {0.5, 0.1, 0.0, 0.0, 0.0, 0.0, 0.0, 0.1, 0.2, 0.3, 1.0}, where 0.5 is the rescaling factor for age 0 individuals and 1.0 is the rescaling factor for age 10 individuals. In other words, these values represent a survival rate multiplier for each age (i.e., age 0 individuals have a relative reduction in survival probability of 0.5, whereas age 1 individuals have a relative reduction of 0.1, and all individuals die after age 10). The resulting survivorship curve in our model is shown in Figure S3. Together, these life history parameters yielded a generation interval of 3 years, consistent with those observed in other felid species, which typically range from ∼2-5 years (Cho et al., 2013; Saremi et al., 2019; Wang et al., 2022). As caracal generation time estimates do not exist, we used this model-based generation time to inform our previously described demographic inference.

### Genomic parameters

We set the genomic parameters of our simulation with the aim of modelling deleterious mutations occurring as nonsynonymous mutations in coding regions. As informed by the domestic cat reference genome (Buckley et al., 2020), we modelled 18 autosomes each with 1100 genes of length 1670bp (19,800 total genes), yielding a total sequence length of ∼33 Mb. We assumed a recombination rate of 1e-6 crossovers per site per generation between genes and 0.5 between chromosomes, with no recombination occurring within genes. We chose this recombination rate to be high enough to avoid elevated linkage between genes though low enough to allow for long ROH>1Mb to accumulate under close inbreeding as a means for measuring F_ROH_ in the simulated population. Within each gene, we modelled deleterious mutations occurring at a rate of 8.6e-9*2.31/3.31=6e-9 mutations per site per generation, where 8.6e-9 is an estimate of mutation rate from a trio-based study in domestic cats (Wang et al., 2022) and 2.31/3.31 is the fraction of coding mutations that are assumed to be nonsynonymous (Huber, Kim, Marsden, & Lohmueller, 2017). Together, these parameters yield a genomic deleterious mutation rate of *U*=0.4 (2*6e-9*18*1100*1670). For computational efficiency, we did not model neutral (synonymous) mutations, as these mutations do not contribute to fitness but do greatly add to the computational overhead of a simulation.

Selection coefficients (*s*) for new deleterious mutations were drawn from a mixture distribution described in Kyriazis et al. (2023) based on a gamma distribution inferred for humans by Kim, et al. (2017). Specifically, we assumed that 99.5% of mutations were drawn from a gamma distribution with a mean *s*=-0.0131 and shape parameter = 0.186, with the other 0.5% of mutations being recessive lethal with *s*=-1.0, as informed by Wade et al. (2023). Dominance coefficients (*h*) for deleterious mutations were set following Kyriazis et al. (2023) as: *h*=0.45 for weakly deleterious mutations (*s*>-0.001), *h*=0.2 for moderately deleterious mutations (-0.001>=*s*>-0.01), *h*=0.05 for strongly deleterious mutations (-0.01>*s*=>-0.001), and *h*=0.0 for lethal and semi-lethal mutations (-1.0<=*s*<-0.1). Note that the mean *h* of new deleterious mutations in this model is 0.28.

### Demographic parameters

The demographic parameters (i.e., carrying capacities, divergence times, and migration rates) of our simulation were set using the two-epoch divergence model estimated above. This model includes an ancestral effective population size (*N_anc_*) of 35,265; effective population size in the Cape Peninsula population (*N_CP_*) of 28; effective population size for the Greater Cape Town population (*N_GCT_*) of 92; and migration fraction of 0.046 when assuming a divergence time of 25 generations (or 75 years, assuming a generation time of 3 years). To convert these effective population size parameters to carrying capacities (which determine the maximum potential census size of the simulated population), we first determined the ratio between *N_e_* and *N* in our model by estimating *N_e_* from neutral heterozygosity at equilibrium (*N_e_* = *π*/(4\**μ*)) for an arbitrary *K* (note that under a neutral model with no deleterious mutations, *N*≈*K*), yielding a *N_e_*/*N* ratio of ∼0.49. We also determined the mean equilibrium absolute fitness in the presence of deleterious mutations in our model to be 0.94, which represents the average ratio between the simulated census population size and carrying capacity (*N*/*K*). Dividing our *N_e_* estimates by these two ratios (*N_e_*/(0.49*0.94)) yields *K_anc_*=∼76,000, *K_CP_*=∼60, and *K_GCT_*=∼200. This estimate of a carrying capacity for the Cape Peninsula population of 60 notably agrees well with the census size estimate of ∼48 individuals (Avenant et al., 2016; Leighton et al., 2022).

The symmetric migration rate estimated from our demographic inference of 0.046 (see below) represents a migration fraction for each population (i.e., each generation, 4.6% of parent chromosomes are drawn from the other population). For the Cape Peninsula population, this translates to an effective migration rate of 0.046*28=∼1.3 effective migrants per generation, or ∼0.43 effective migrants per year. In the SLiM nonWF model, migration is enforced by moving migrants between populations each year, where the number of actual migrants moved is not identical to the effective number of migrants, as there is no guarantee that a migrant will reproduce after translocation. Thus, the relationship between the actual migration rate and effective migration rate is not necessarily straightforward (see Petkova et al. (2016) for further discussion). With these complexities in mind, we assumed symmetric migration rates of 1 migrant per generation from 1940-1990 and 0.5 migrants per generation from 1991-2025, assuming that the actual number of migrants each year was Poisson distributed. This downward shift of the migration rate was assumed to model the effects of the growing population density of Cape Town, which likely had the effect of reducing migration corridors for caracals. Note that these actual migration rates were chosen to be somewhat higher than our estimate of the effective migration rate, given the fact that not all migrants will survive and reproduce. To check that these migration rates are sensible, we compared the estimated F_ST_ in our genomic dataset to that predicted by our model (see Results).

### Modelling stochastic anthropogenic mortality

In addition to inbreeding depression, a major threat to the Cape Peninsula caracal population is anthropogenic mortality. Specifically, caracals are commonly killed by collisions with vehicles, poison, poaching, and fires (Leighton et al., 2022; Parker et al., 2023; Serieys et al., 2019; Viljoen et al., 2020). To incorporate this into our model, we enforced a probability of stochastic anthropogenic mortality (*p_mort_*) where individuals in the Cape Peninsula population were randomly killed each year. To set *p_mort_*, we determined the average number of mortalities per year during the period 2015-2022 as being 7.9 (range 5-13). Note that this is likely to be an underestimate, as some mortalities may go undetected. Thus, based on the assumed carrying capacity of 60, we set *p_mort_*=0.15 (0.15*60=9 mortalities per year on average). Given that mortality rates were likely lower in the past, when population densities were lower, we assumed *p_mort_*=0.05 from 1940-1990, *p_mort_*=0.1 from 1990-2010, and *p_mort_*=0.15 from 2010-2070 by default. We simulated mortality each year as a Bernoulli trial for each individual.

### Simulating future migration and mortality scenarios

Following decline and divergence, we modelled population dynamics as described above from 1940-2025, and from 2025-2070 we explored various hypothetical migration and mortality scenarios (Table 1). These scenarios consist of (1) models where the natural stochastic rate of migration varied, (2) models where additional migrants were translocated to the Cape Peninsula to initiate a genetic rescue, and (3) models where the stochastic anthropogenic mortality rate was varied.

**Table 1:** Description of parameters that were varied in simulations including resulting extinction rates and mean *N* in 2070. Parameters include the ancestral carrying capacity (*K_anc_*_),_ the natural stochastic migration rate in # of individuals per year (mig_rate), the annual probability of anthropogenic mortality (*p_mort_*), the number of migrants sent for genetic rescue every 5 years (#rescue), the genomic deleterious mutation rate (*U*), and the mean dominance coefficient (mean *h*). Percent extinct and mean *N* in 2070 are reported for the Cape Peninsula population.

| name | $K_{anc}$ | mig_rate | $p_{mort}$ | #rescue | $U$ | mean $h$ | % extinct | $N$ in 2070 |
| --- | --- | --- | --- | --- | --- | --- | --- | --- |
| baseline model | 76000 | 0.5 | 0.15 | 0 | 0.4 | 0.28 | 2 | 41.8 |
| **Migration and mortality scenarios** |  |  |  |  |  |  |  |  |
| *mig0.0* | 76000 | 0.0 | 0.15 | 0 | 0.4 | 0.28 | 11 | 32.1 |
| *mig2.0* | 76000 | 2 | 0.15 | 0 | 0.4 | 0.28 | 2 | 42.3 |
| *rescue* | 76000 | 0.5 | 0.15 | 5 from GCT | 0.4 | 0.28 | 0 | 44.8 |
| *rescue_div* | 76000 | 0.5 | 0.15 | 5 from CK | 0.4 | 0.28 | 0 | 45.5 |
| *pmort0.1* | 76000 | 0.5 | 0.1 | 0 | 0.4 | 0.28 | 1 | 47.0 |
| *pmort0.2* | 76000 | 0.5 | 0.2 | 0 | 0.4 | 0.28 | 24 | 24.0 |
| **Genomic and demographic scenarios** |  |  |  |  |  |  |  |  |
| $U=0.3$ | 76000 | 0.5 | 0.15 | 0 | 0.3 | 0.28 | 0 | 45.6 |
| $U=0.5$ | 76000 | 0.5 | 0.15 | 0 | 0.5 | 0.28 | 19 | 30.3 |
| $h=0.35$ | 76000 | 0.5 | 0.15 | 0 | 0.4 | 0.35 | 0 | 44.4 |
| $Ka=7600$ | 7600 | 0.5 | 0.15 | 0 | 0.4 | 0.28 | 0 | 42.9 |

To facilitate interpretation of these results, we determined a ‘baseline model’ as being the case where stochastic migration rates and mortality rates remain the same in the future (0.5 migrants per year, 0.15 mortality rate), with no genetic rescue. For all scenarios, we varied only one of these model elements relative to the baseline model. Specifically, we explored scenarios where natural migration rates either decreased to 0 migrants per year (*0.0mig*) or increased to 2 migrants per year (*2.0mig*; Table 1) from 2025-2070. To explore the impact of genetic rescue, we implemented a scenario where five migrants were translocated every five years from the Greater Cape Town population to the Cape Peninsula population from 2025-2070 (*rescue*). To explore the potential impact of using a more divergent source population for genetic rescue (*rescue_div*), such as the Central Karoo or Namaqualand population, we also explored a genetic rescue scenario where we split off a third population in 1940 with *K*=200 and no migration connectivity with the Cape Peninsula. We then used this more divergent population as a source for sending five migrants every five years. Note that F_ST_ between the Cape Peninsula and this more divergent source population was 0.09, similar to the estimated F_ST_ between the Cape Peninsula and Central Karoo (F_ST=_0.10), as well as Cape Peninsula and Namaqualand (F_ST_=0.11). Finally, to explore the impact of varying the stochastic anthropogenic mortality rate (*p_mort_*), we ran models where this parameter either decreased to 0.1 (*pmort0.1*) or increased to 0.2 (*pmort0.2*) from 2025-2070.

### Sensitivity analyses of genomic and demographic parameters

To determine the extent to which model projections were influenced by our assumed genomic and demographic parameters, we ran sensitivity analyses where we varied the genomic deleterious mutation rate, distribution of dominance coefficients, and historical population size. Here, we employed the same ‘baseline model’ described above as a point of reference, which included a genomic deleterious mutation rate of *U*=0.4, mean *h*=0.28, and *K_anc_*=76,000 (Table 1). For all sensitivity analyses, we assumed a future migration rate of 0.5 migrants per year, stochastic mortality rate of 0.15, and no genetic rescue.

To explore the impact of varying the genomic deleterious mutation rate, we ran simulations with an increased mutation rate (*U*=0.5) as well as a decreased mutation rate (*U*=0.3; Table 1). We implemented this varying mutation rate by changing the length of coding sequence in our model from the default of 33Mb to a larger (41.6Mb) and smaller (24.8Mb) size while keeping the per site mutation rate constant. Given uncertainty about dominance parameters in mammals (Kyriazis et al., 2023), we next ran simulations with a somewhat less recessive distribution of *h* with: *h*=0.5 for weakly deleterious mutations (*s*>-0.001), *h*=0.35 for moderately deleterious mutations (-0.001>=*s*>-0.01), *h*=0.1 for strongly deleterious mutations (-0.01>*s*=>-0.001), and *h*=0.01 for semi-lethal mutations (-1.0<*s*<-0.1), and *h*=0.0 for lethal mutations (*s*=-1.0). Note that the overall mean *h* of deleterious mutations in this model is 0.35 (Table 1). Finally, we also sought to determine the degree to which genetic load, inbreeding load, and extinction risk in our model depended on the large ancestral population size (*K_anc_*=76,000). To explore the effects of assuming a much smaller ancestral population size, we ran simulations with *K_anc_*=7,600.

### Model implementation and output

For each scenario described above, we ran 100 simulation replicates. We ran burn-ins for 5,000 years at *K_anc_*, followed by divergence and decline to *K_CP_* and *K_GCT_*. This burn-in duration was chosen to be long enough for the simulated inbreeding load to reach equilibrium (Fig. S4) though short enough to maintain computational feasibility (note that each replicate takes ∼20-50 hours and up to 8G of memory on our compute cluster). During each simulation replicate, we kept track of several quantities. These include the simulated population size, mean levels of inbreeding (F_ROH_ for ROH>1Mb), mean F_ST_ between the Cape Peninsula and Greater Cape Town populations, as well as the simulated genetic load and inbreeding load. Note that genetic load is measured here as 1-mean absolute fitness, whereas inbreeding load is measured as the ‘diploid number of lethal equivalents’, or the summed *s* of heterozygous recessive deleterious mutations (Hedrick & Garcia-Dorado, 2016; Kyriazis et al., 2023; Morton, Crow, & Muller, 1956). In other words, the genetic load represents the realized loss of fitness in the simulated population (i.e., realized load) whereas the inbreeding load measures the quantity of recessive deleterious variation that is masked as heterozygotes (i.e., masked load; Bertorelle et al., 2022). Here, we measure inbreeding load for annual survival as well as survival to 3 years (the assumed generation interval), as this quantity is often reported either for juvenile survival or survival to sexual maturity (Nietlisbach, Muff, Reid, Whitlock, & Keller, 2019). These quantities were outputted every year from a sample of 30 individuals.

## Results

### Sampling & population structure

We sequenced 31 caracal genomes to high coverage (mean coverage = 31.5x, range 25.0-43.2; Table S1). Raw reads mapped well to the domestic cat genome, with a mean mapping rate of 98.7% (Table S1). Principal component analysis of these 31 genomes depicted notable structure among sampled populations (Fig. 1b). Specifically, all four populations clustered by geographic origin when plotting the first two principal components, with the exception of CM32 (Fig. 1b). This individual was sampled in the Cape Peninsula but grouped with the Greater Cape Town population in the PCA. Moreover, patterns of separation in the PCA roughly reflected geography, with the Greater Cape Town individuals being in the center of the PCA and the three other populations branching out (Fig. 1b). Hierarchical clustering based on identity-by-state similarly found evidence for population structure, with individuals generally clustering together by geographic origin (Fig. S5). Additionally, estimates of pairwise kinship coefficients found evidence of high relatedness among Cape Peninsula samples, with three individuals having kinship coefficients >0.1 (Fig. S6), including a high kinship coefficient of 0.20 between C23 and CM32 (note that the expected kinship coefficient for parent-offspring is 0.25). This finding, together with the age and sampling dates of these individuals, suggests that C23 may be the mother of CM32 and that the father of CM32 is likely a migrant from the Greater Cape Town population. Finally, estimates of genome-wide F_ST_ suggest that, although population structure is evident, overall differentiation is relatively low between the Cape Peninsula population and Greater Cape Town population, as evidenced by a pairwise F_ST_ estimate of 0.053, compared to an overall estimate of F_ST_ across all four populations of 0.085.

### Genetic diversity & inbreeding

We next sought to determine whether isolation in the Cape Peninsula has resulted in decreased levels of genetic diversity or increased levels of inbreeding. We found that heterozygosity is significantly reduced in the Cape Peninsula relative to the three other sampled populations (Fig. S7, S8), and that this reduction in genetic diversity has been driven largely by recent inbreeding (Fig. 2). Specifically, the mean heterozygosity across Cape Peninsula individuals was found to be *π*=9.0e-4 pairwise differences per site, an 11.5% reduction relative to that observed in the Greater Cape Town, Central Karoo, and Namaqualand populations (mean *π*=1.1e-3; Fig. S7). Across all samples, we observe a strong correlation between individual heterozygosity and F_ROH_ for runs of homozygosity >1Mb in length (Fig. 2a), suggesting that variation in genetic diversity across populations and individuals is driven largely by inbreeding. Runs of homozygosity were present across nearly all sampled individuals, most notably in the Cape Peninsula (mean F_ROH_=0.20), but also within the Greater Cape Town, Central Karoo, and Namaqualand populations (mean F_ROH_=0.12; Fig. 2). This higher level of inbreeding in the Cape Peninsula was driven largely by an abundance of long ROH>10Mb, which were generally absent from other sampled populations (Fig. 2b, d). For instance, several ROH were found to encompass more than 30Mb in length (Fig. 2b), at times covering roughly half of an individual chromosome (Fig. 2d, S9). However, a notable exception to this was found for CM32, an individual that had among the highest heterozygosity (*π*=1.2e-4) and lowest inbreeding coefficient (F_ROH_=0.059) of any individual in our dataset (Fig. 2a) despite originating from the Cape Peninsula. Although the putative mother of CM32 was highly inbred (F_ROH_=0.21), the divergent set of haplotypes provided by the putative migrant father likely served to complement and mask this inbreeding in CM32.

**Figure 2:**
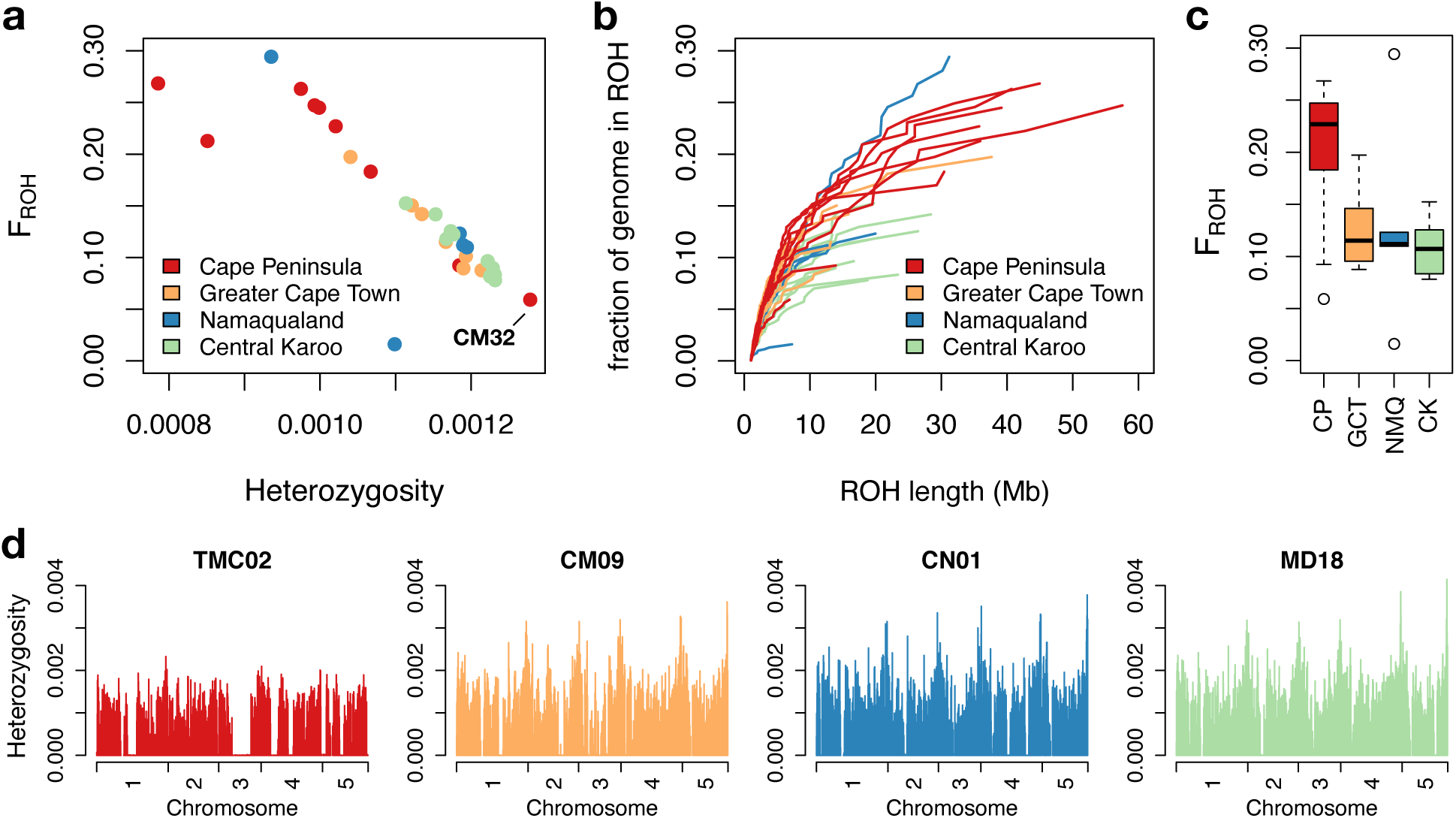
Patterns of genetic diversity and inbreeding. (a) Low genetic diversity in the Cape Peninsula caracal population is driven by high levels of inbreeding. Scatterplot depicts relationship between heterozygosity and the inbreeding coefficient for each individual, as measured by F_ROH>1Mb_. Note that Cape Peninsula individuals generally have much lower heterozygosity and higher levels of inbreeding, with the exception of CM32. (b) High levels of inbreeding in the Cape Peninsula driven primarily by long ROH >10Mb. Plot shows the cumulative proportion of individual genomes covered by ROH of increasing length. (c) Boxplots showing the distribution of F_ROH>1Mb_ in each population. See Figure S7 for boxplots of heterozygosity for each population. (d) Plots of heterozygosity in 1Mb sliding windows for one representative individual from each population. To facilitate visualization, results shown for only the first five chromosomes. Note the abundance of long ROH present in TMC02 from the Cape Peninsula, which are largely absent from other individuals. See Figure S9 for plots of all individuals.

The abundant long ROH present in the Cape Peninsula population suggest that reduced genetic diversity in this population has been driven largely by recent inbreeding, possibly due to the isolation of the peninsula during the expansion of Cape Town over the past several decades. To more precisely date the age of observed ROH, we used an approach for modelling the decay of ROH (Browning, 2008; Thompson, 2013), finding a mean age for ROH>1Mb of 10.8 generations and a mean age for ROH>10Mb of 2.4 generations (Table S2). Assuming a generation time of 3 years, these translate to 32.4 years and 7.2 years, respectively. These results suggest that inbreeding in the Cape Peninsula is a relatively recent phenomenon that has been driven largely by population isolation over the past several decades. However, the presence of inbreeding across other sampled populations also suggests that population declines and/or isolation may be occurring across the caracal’s range in South Africa. For instance, in the Central Karoo population, we similarly estimate a mean ROH>1Mb age of 14.8 generations (44.3 years) as well as a mean ROH>10Mb age of 3.1 generations (9.4 years; Table S2). Thus, recent inbreeding appears to be present in both urban and non-urban caracal populations in South Africa, though is most prevalent in the isolated Cape Peninsula.

### Inferring ancestral and contemporary effective population sizes and migration rates

To estimate historical and contemporary effective population sizes and migration rates for the Cape Peninsula and Greater Cape Town populations, we fitted a divergence model to the SFS using ∂a∂i (Gutenkunst et al., 2009). We observed good convergence and fit to the empirical data for a model with a large *N_anc_*=35,265, splitting into two much smaller populations with *N_CP_*=28 and *N_GCT_*=92 when assuming a divergence time of 25 generations (Figs. 3, S11-S12). In this model, we also estimate a symmetric migration fraction of 0.046, suggesting a rate of effective migration into the Cape Peninsula of 1.29 migrants per generation over the past 75 years, or 0.43 migrants per year.

**Figure 3:**
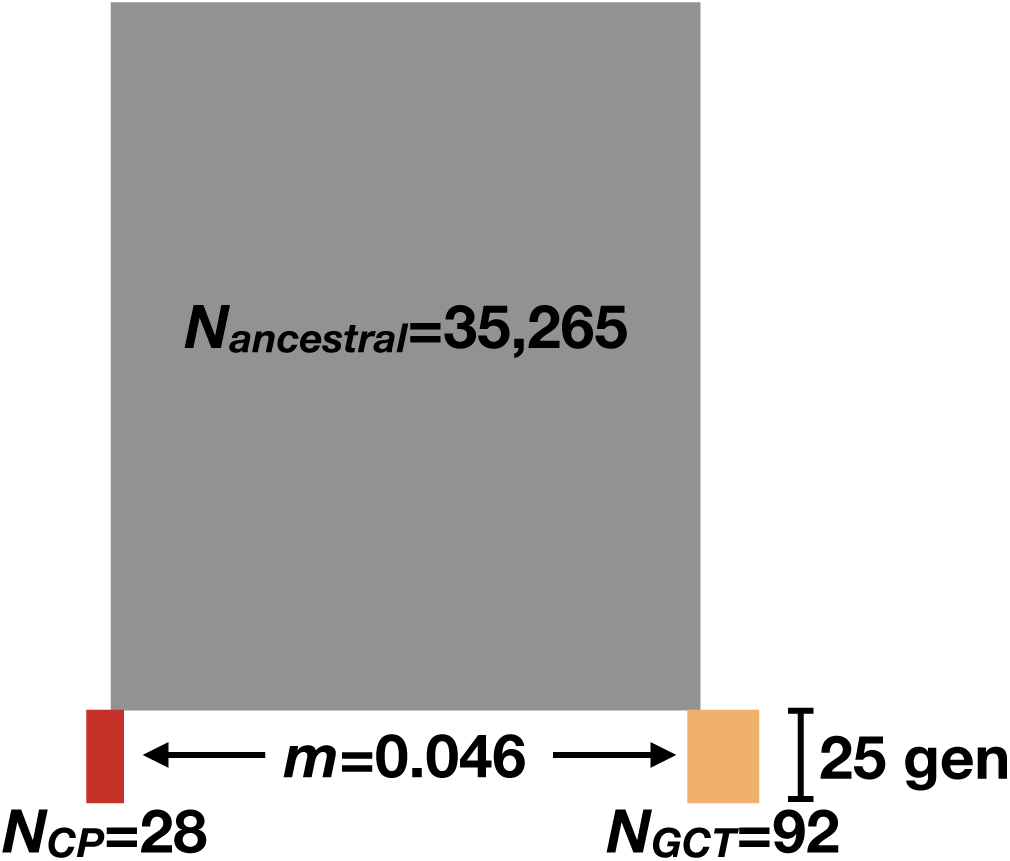
Divergence model estimated using ∂a∂i for the Cape Peninsula and Greater Cape Town populations. Depicted are the estimated effective population sizes and symmetric migration rate between the Cape Peninsula and Greater Cape Town populations. Note that this model assumes a divergence time of 25 generations (see Methods). See Figures S11 and S12 for plots showing the fit of the two-dimensional and marginal SFS.

Importantly, the large estimated *N_anc_*=35,265 in our model likely reflects the broader caracal population in South Africa, which was historically large and connected by a high degree of migration. However, the consequence of population fragmentation and declines is that our model estimates very small recent effective population sizes in the Greater Cape Town and Cape Peninsula, as these populations are no longer highly connected by migration to the broader population in South Africa.

### Simulation results under the baseline model

Our eco-evolutionary simulation results projecting population dynamics in the Cape Peninsula into the future highlight the complex interplay between migration, anthropogenic mortality, deleterious variation parameters, and population viability. To summarize these results, we will first describe results from our ‘baseline model’ in detail, followed by results while varying migration, mortality, genomic, and demographic parameters.

In our baseline model with a future migration rate of 0.5 migrants per year, a future anthropogenic mortality rate (*p_mort_*) of 0.15, *U*=0.4, *K_anc_*=76,000, and mean *h*=0.28 (Table 1), we observe moderate declines in population size (Fig. 4). Specifically, the average census population size in our model declined from 50.3 in 1940, to 43.3 in 2020, to 41.8 in 2070 (Table 1; Fig 4). Although only 2% of replicates are predicted to go extinct by 2070 (Table 1), 29% are predicted to decline to *N*<40. These declines are driven, in part, by increasing levels of inbreeding: mean F_ROH_ in the Cape Peninsula increased from F_ROH_=0.003 in 1940, to F_ROH_=0.18 in 2020, and F_ROH_=0.25 in 2070 (Fig. 4). Similar trends are observed for F_ST_ between the Cape Peninsula and Greater Cape Town populations, which increased from F_ST_=0.011 in 1940, to F_ST_=0.054 in 2020, and F_ST_=0.061 in 2070 (Fig. 4). The projected mean F_ROH_=0.18 and F_ST_=0.054 in 2020 notably agrees with the empirical estimates of F_ROH_=0.20 and F_ST_=0.053 (Fig. 4). This good agreement between model predictions and empirical estimates suggests that our demographic parameters are well calibrated, serving as an additional source of evidence to validate our model.

**Figure 4:**
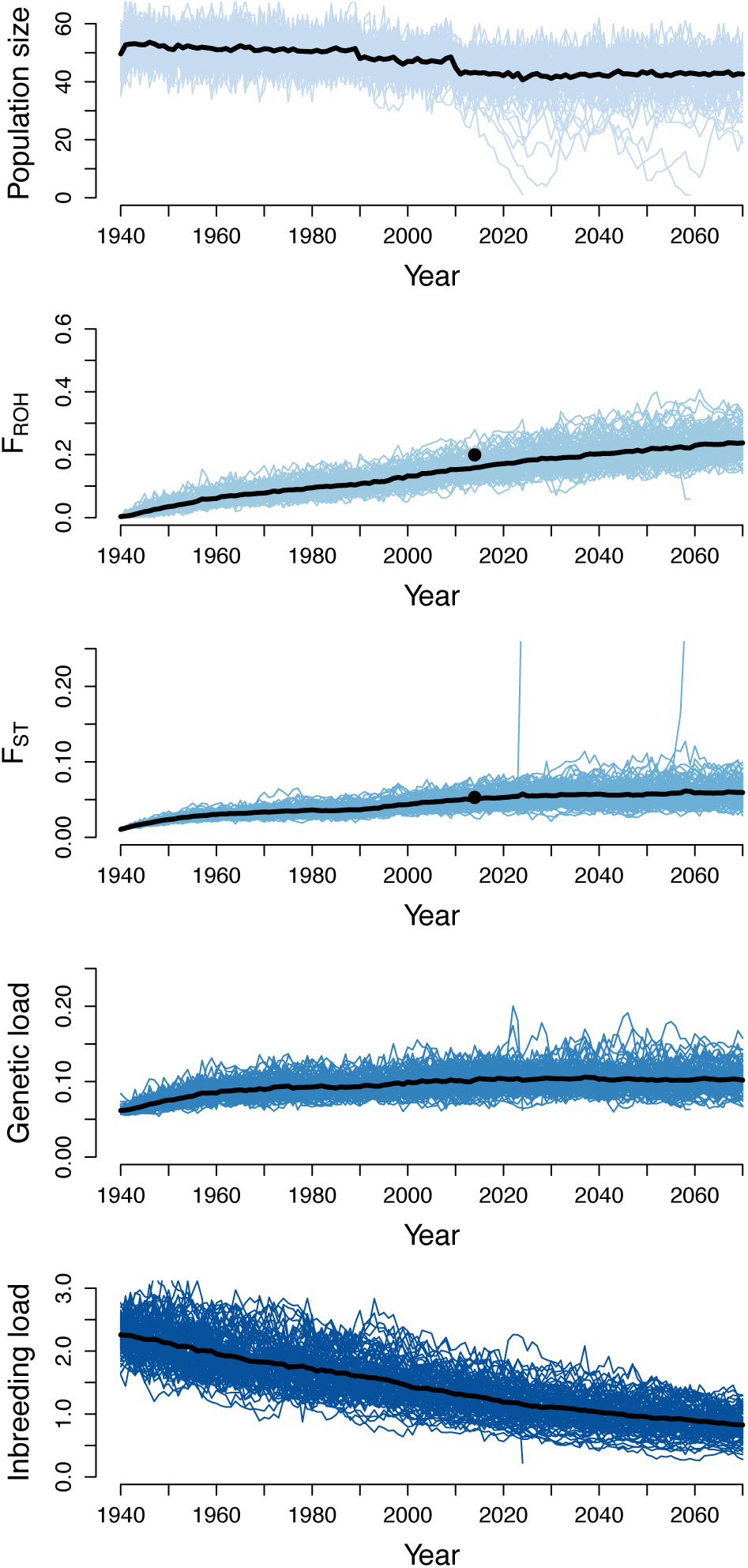
Eco-evolutionary simulation results for the Cape Peninsula population under the ‘baseline model’ where migration rates remain the same in the future (0.5 migrants per year). This baseline model further assumes *K_anc_*=76,000, a stochastic anthropogenic mortality rate of 0.15 in the future, and a genomic deleterious mutation rate of *U*=0.4 (see main text for more details). For this model, we show the simulated population size, mean inbreeding coefficient (F_ROH_), fixation coefficient (F_ST_) between the Cape Peninsula and Greater Cape Town population, mean genetic load, and mean inbreeding load for annual survival from 1940-2070. Each panel shows trajectories for all 100 simulation replicates as well as the mean trajectory in black. The dots in the F_ROH_ and F_ST_ panels indicate the empirical genomic estimates.

As a consequence of increased isolation and inbreeding in the simulated Cape Peninsula population, our model predicts an accumulation of genetic load as well as a decrease (or purging) of the inbreeding load (Fig. 4). Specifically, genetic load increased from 0.062 in 1940 to 0.10 in 2020 and finally to 0.11 in 2070 (Fig. 4). By contrast, the simulated inbreeding load decreased from 2*B*=2.29 in 1940 to 2*B*=1.2 in 2020 and finally to 2*B*=0.82 in 2070 (Fig. 4). Note that inbreeding load is measured here as the diploid number of lethal equivalents for annual survival; when measured over the generation time of caracals (3 years) our model predicts inbreeding loads of 2*B*=4.67, 2*B*=2.70, and 2*B*=1.99 in 1940, 2020, and 2070, respectively. Taken together, these results suggest that, under our baseline model parameters where present-day rates of migration and anthropogenic mortality remain constant in the future, further population declines, increases in inbreeding, population divergence, and genetic load are likely to occur in the Cape Peninsula. However, population extinction appears unlikely to occur within the next ∼50 years under a ‘business as usual’ scenario. This may be enabled to some degree by a purging of the inbreeding load as the Cape Peninsula population becomes more inbred, as well as the beneficial effects of a migration rate that, though low, remains above zero.

### Simulation results while varying migration and mortality parameters

We next sought to determine how our model projections may be impacted by varying the rate of future migration (either naturally or via genetic rescue) and rate of stochastic anthropogenic mortality. When assuming no migration from 2025-2070 (*0.0mig*), our model predicts a substantially reduced average population size in 2070 of 32.1, with 11% of replicates declining to extinction (Table 1; Fig. 5). Moreover, this model also predicts greatly elevated levels of inbreeding, population differentiation, and genetic load compared to the baseline model (Fig. 5). By contrast, if migration rates increase to 2 migrants per year from 2025-2070 (*2.0mig*), the predicted mean population size in 2070 remains at 42.3 individuals and just 2% of replicates are predicted to go extinct (Table 1; Fig. 5). In this scenario, predicted F_ROH_, F_ST_, and genetic load also remain relatively low in 2070, though much less of the inbreeding load is purged (Fig. 5).

**Figure 5:**
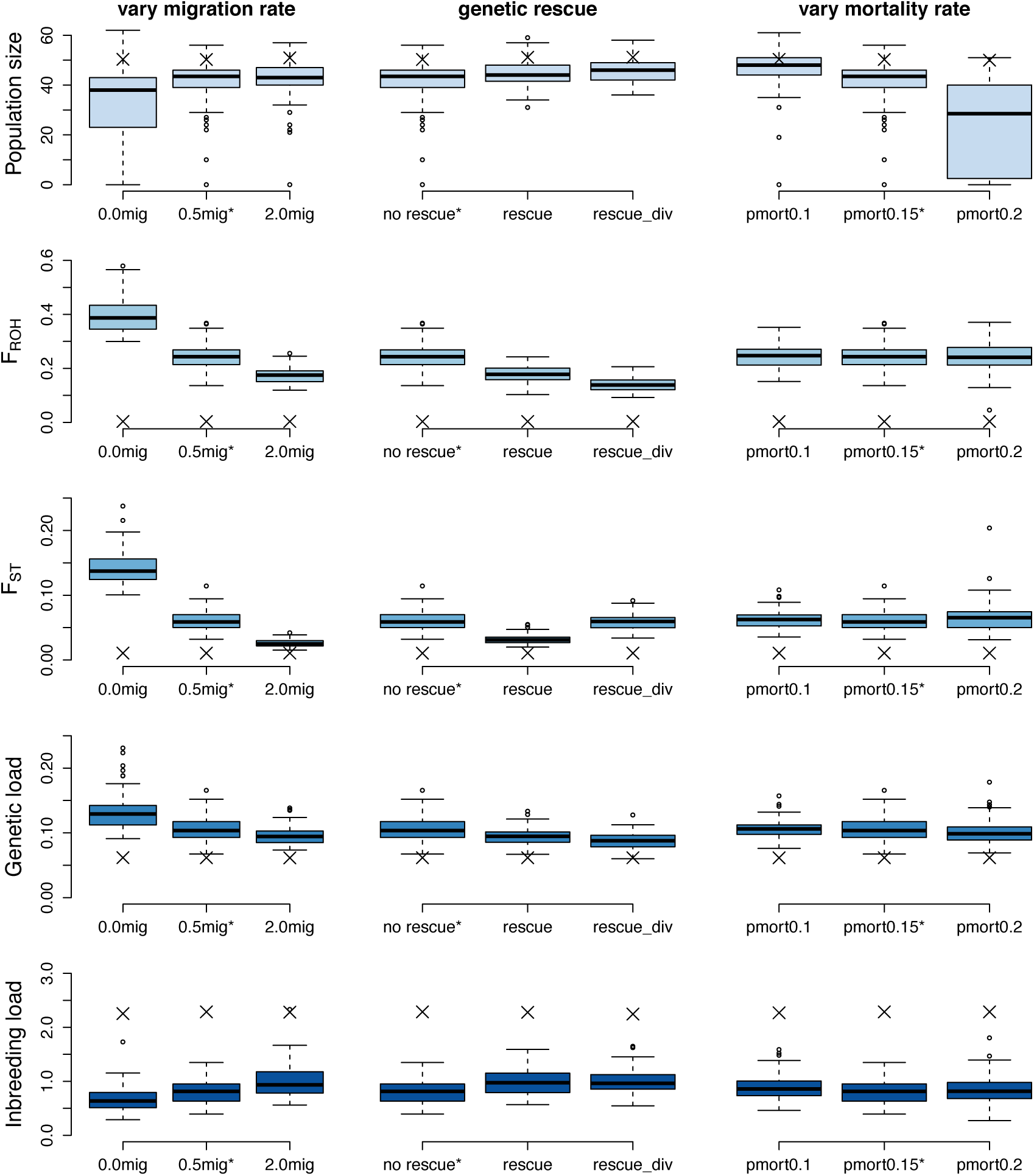
Simulation projections in 2070 under various future migration and mortality scenarios. Boxplots show the distribution of each simulated quantity in the Cape Peninsula population in 2070, with the average starting value for each quantity in 1940 shown with an ‘X’. Quantities shown include the simulated population size, mean inbreeding coefficient (F_ROH_), fixation coefficient (F_ST_) between the Cape Peninsula and Greater Cape Town populations, mean genetic load, and mean inbreeding load. The distribution for each of these quantities is shown for scenarios with varying future migration rates (left column), genetic rescue (middle column), and varying stochastic anthropogenic mortality rates (right column). Note that the ‘baseline model’ shown in Figure 4 is denoted with an asterisk. See Table 1 for extinction rates and mean population size in 2070 for each scenario.

Although facilitating natural migration may represent the ideal approach for maintaining gene flow to the Cape Peninsula population, implementing this is likely not achievable due to the extent of dense development near the Cape Peninsula (Fig. 1, S1). Given this reality, we explored two genetic rescue scenarios in which five migrants were translocated to the Cape Peninsula every five years during 2025-2070 either from the Greater Cape Town population (*rescue*) or a more divergent population such as the Central Karoo or Namaqualand (*rescue_div*). In both the *rescue* and *rescue_div* scenario, our model predicts notably high population sizes in 2070 of 44.8 and 45.5 individuals, respectively, along with decreased levels of inbreeding, population divergence, and genetic load (Table 1; Fig. 5). Interestingly, however, we observed the lowest levels of inbreeding and genetic load when sourcing migrants from a more divergent population (Fig. 5), perhaps due to the divergent source population providing haplotypes carrying distinct alleles at many recessive deleterious loci. These distinct haplotypes can better mask high-frequency recessive deleterious alleles in the Cape Town population, leading to greater heterosis. Nevertheless, both genetic rescue scenarios were sufficient to ensure that population sizes remained relatively high and no replicates went extinct by 2070 (Table 1; Fig. 5).

We next explored the effect of varying the stochastic anthropogenic mortality rate (*p_mort_*), which was set to 0.15 in the baseline model. When decreasing this rate to 0.10 from 2025-2070 (*pmort0.1*), we observe the largest mean population size in 2070 of any simulated scenario of 47.0, with 1% of replicates going extinct (Table 1; Fig. 5). By contrast, an increase in the mortality rate to 0.20 (*pmort0.2*) resulted in a greatly reduced average population size in 2070 of 24.0, with 24% of replicates going extinct (Table 1; Fig. 5). Notably, varying the anthropogenic mortality rate did not have any noticeable impacts on the predicted F_ROH_, F_ST_, genetic load, or inbreeding load in the Cape Peninsula population in 2070 (Fig. 5).

### Simulation results while varying genomic and demographic parameters

All modelling results so far assume a genomic deleterious mutation rate of *U*=0.4 and average dominance coefficient for deleterious mutations of *h*=0.28. Given uncertainty in these parameters (Kyriazis et al., 2023), we next aimed to explore how our model projections may change when varying the deleterious mutation rate and distribution of dominance coefficients. When modelling a higher deleterious mutation rate of *U*=0.5, we observe a decreased mean population size in 2070 of 30.3 with 19% of replicates going extinct (Table 1). This much lower predicted population size is due to the simulated population having a greatly elevated genetic load and inbreeding load (Fig. 6), a direct consequence of the higher rate of deleterious mutation. When decreasing the genomic deleterious mutation rate to *U*=0.3, we observe the opposite outcome: the average population size in 2070 is 45.6 individuals, with no replicates going extinct (Table 1). As expected, the predicted inbreeding load and genetic load is greatly diminished in this model (Fig. 6), resulting in a lessened impact of deleterious mutations on population dynamics. The effects of modelling a somewhat less recessive distribution of dominance coefficients (mean *h*=0.35) similarly leads to a reduced influence of deleterious mutations on population dynamics. Specifically, the average predicted population size in 2070 under this model was 44.4 with no replicates going extinct (Table 1), a consequence of a lowered inbreeding load when mutations are less recessive, translating to a reduced genetic load after the population becomes inbred.

**Figure 6:**
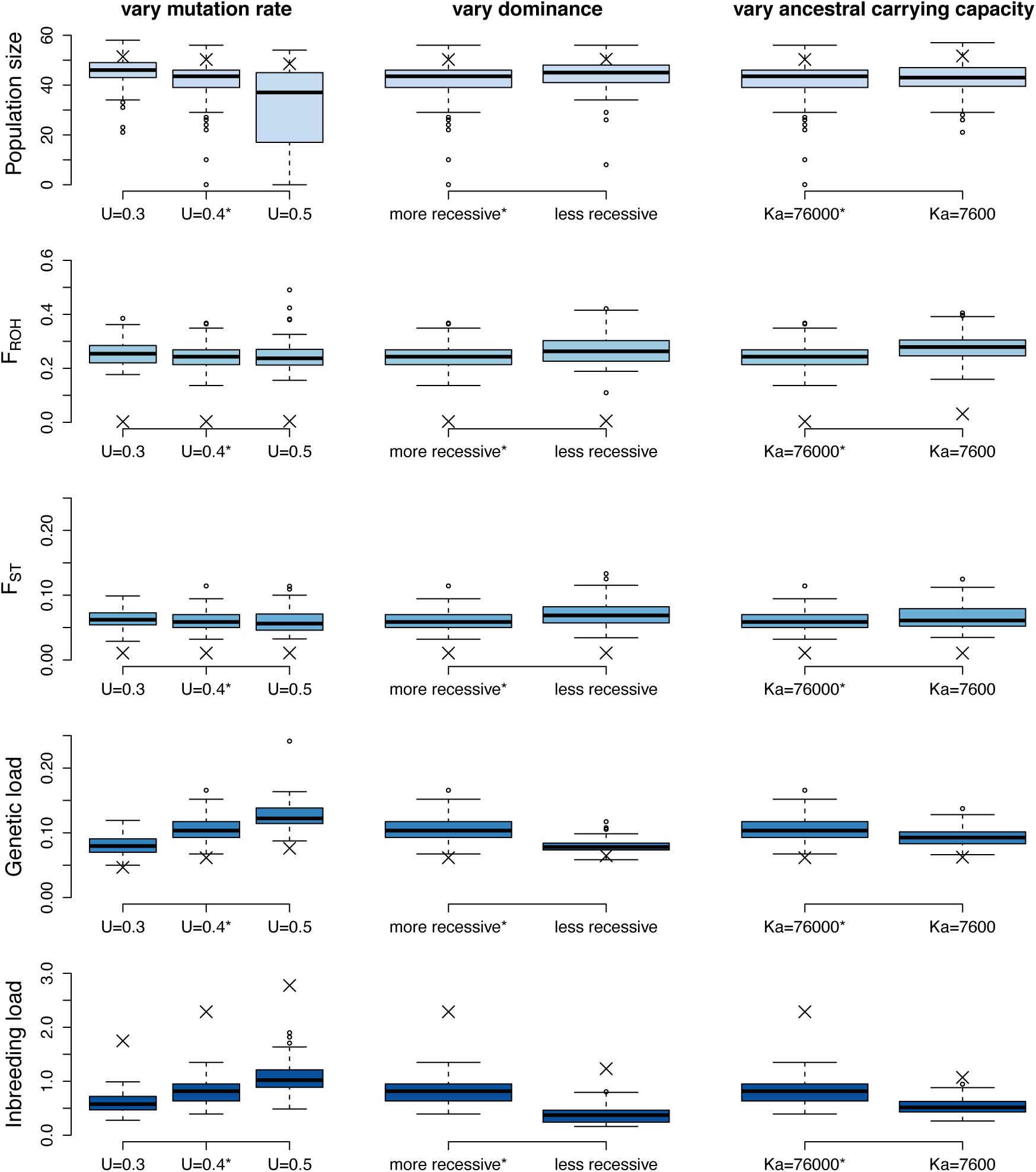
Simulation projections in 2070 under various genomic and demographic parameters. Boxplots show the distribution of each simulated quantity in the Cape Peninsula in 2070, with the average starting value for each quantity in 1940 shown with an ‘X’. Quantities shown include the simulated population size, mean inbreeding coefficient (F_ROH_), fixation coefficient (F_ST_) between the Cape Peninsula and Greater Cape Town populations, mean genetic load, and mean inbreeding load. The distribution for each of these quantities is shown for scenarios with varying deleterious mutation rates (left column), dominance distributions (middle column), and historical population sizes (right column). Note that the ‘baseline model’ shown in Figure 4 is denoted with an asterisk. See Table 1 for extinction rates and mean population size in 2070 for each scenario.

One final scenario we explored was a model where the ancestral carrying capacity was reduced by an order of magnitude to *K_anc_*=7,600. The motivation for this scenario was to examine the extent to which our model projections depend on the large historical population size we estimate for caracals (Fig. 3), given that many recent studies have suggested that ancestral population size may play an important role in determining extinction risk in small populations (Angeloni et al., 2011; Kyriazis et al., 2021; Pekkala, Knott, Kotiaho, & Puurtinen, 2012; Robinson et al., 2022, 2023, 2019; Wilder et al., 2023). When modelling a reduced *K_anc_*, we observe only modest effects on population dynamics, with a predicted population size in 2070 of 42.9 (Table 1; Fig. 6). However, the predicted inbreeding load and genetic loads are greatly lowered when *K_anc_*=7,600, as expected (Fig. 6).

## Discussion

In this study, we have integrated whole genome sequencing data with computational simulation modelling to investigate the impact of habitat fragmentation and reduced gene flow on population viability. We estimate a reduced rate of migration into the Cape Peninsula, contributing to elevated population structure (Fig. 1b, Fig. S5). This reduced migration, together with the small effective population size in the Cape Peninsula (*N_e_*=28; Fig. 3), has led to decreased genetic diversity due to elevated levels of recent inbreeding (Figs. 2, S7-S10). Runs of homozygosity were abundant across individuals sampled from the Cape Peninsula, with a mean F_ROH>1Mb_=0.20 (Fig. 2, S10), nearly as high as that of a full-sibling mating. Much of the homozygosity in the Cape Peninsula appears to have been driven by very recent inbreeding in the form of abundant long ROH>10Mb, which were largely not present in other sampled populations (Fig. 2, S10). The average age for ROH>1Mb in the Cape Peninsula was estimated to be 32.4 years, with an average age of just 7.2 years for ROH>10Mb (Table S2). Notably, these dates are in good agreement with the timing of rapid anthropogenic development in the Cape Flats region starting in the 1980s, perhaps the primary factor driving isolation of the Cape Peninsula (Rebelo et al., 2011). Thus, these results strongly implicate the recent growth of Cape Town as being the primary driver of increased isolation and inbreeding and of reduced genetic diversity in the Cape Peninsula caracal population. Moreover, our finding of abundant long ROH in the Cape Peninsula population suggests that it may already be experiencing reduced fitness due to inbreeding depression. Long ROH are expected to have the most severe effects on fitness, due to arising from recent inbreeding with little opportunity for purging of recessive deleterious variants (Stoffel, Johnston, Pilkington, & Pemberton, 2021; Szpiech et al., 2013). Although we do not have direct measures of fitness in the Cape Peninsula caracal population, work in other natural populations has consistently found a strong link between long ROH and decreased fitness (Stoffel et al., 2021; Swinford et al., 2022), suggesting that such fitness effects may be present in the Cape Peninsula caracal population.

Despite our finding that the Cape Peninsula population appears to be increasingly isolated and inbred, some degree of migration appears to be ongoing. Specifically, we estimate a migration rate of 1.3 effective migrants per generation over the past ∼75 years (Fig. 3), or 0.43 effective migrants per year. Moreover, we directly observe migration in action in our dataset, finding that one individual sampled in the Cape Peninsula (CM32) grouped with individuals from the Greater Cape Town population in our PCA (Fig. 1b). Based on knowledge of this individual and the finding that another Cape Town individual (C23) appears to be the mother of CM32 (Fig. S6), we hypothesize that CM32 is the offspring of an unsampled male migrant. The outcome of this successful migration and reproduction is that CM32 is among the least inbred individuals in our dataset (F_ROH_=0.059), despite her mother being highly inbred (F_ROH_=0.21; Figs. 2a, S9-S10). This is likely due to the putative migrant father providing a distinct set of haplotypes that complemented those of the mother and led to a reduced prevalence of ROHs in their offspring. However, CM32 unfortunately did not survive to reproductive age, as she was killed as a juvenile after being hit by a car. This example is illustrative of the multifarious threats that caracals face in the dense urban environment of Cape Town: even if an individual is able to overcome the challenges of successfully migrating to the Cape Peninsula and reproducing, this alone does not guarantee that their alleles will spread into the broader population.

To model the potential impact of these various threats to persistence in the Cape Peninsula population, we employed the non-Wright-Fisher model in SLiM (Haller & Messer, 2016, 2019, 2023). Specifically, we sought to determine how varying levels of migration and anthropogenic mortality may impact future population dynamics and evaluate whether assisted migration may be warranted. Overall, our simulation results suggest that the Cape Peninsula caracal population is expected to slightly decline under a ‘business-as-usual’ scenario where migration rates and anthropogenic mortality rates do not change in the future (Fig. 4; Table 1). These declines are due to the synergistic effects of anthropogenic mortality as well as an accumulation of genetic load due to increased inbreeding. Specifically, our results demonstrate a strong link between levels of inbreeding, genetic load, and the predicted population size in 2070 (Fig. 5), suggesting that inbreeding depression is a key factor influencing population dynamics. To counteract the effects of inbreeding, we find that migration can serve the important role of providing novel haplotypes that can mask homozygous recessive deleterious mutations and restore fitness. However, despite the increasing levels of inbreeding predicted in our model, extinction risk appears to be relatively low in the near future in most modeled scenarios (Table 1, Figs. 5 & 6). Important exceptions to this are scenarios where future migration is completely halted or where anthropogenic mortality rates further increase, as well as scenarios with a higher genomic deleterious mutation rate (Table 1; Figs. 5 & 6; see below for further discussion). Finally, although the large historical caracal population size appears to have contributed to a high inbreeding load, the effect of this on population dynamics appears to be somewhat less important compared to other elements of our model (Table 1; Fig. 6).

To ameliorate future population decline, our simulations highlight several potential strategies: (1) bolster/maintain the rate of natural gene flow, (2) augment levels of migration by translocating individuals with the aim of performing a genetic rescue, and (3) reduce the level of anthropogenic mortality in the population. In particular, we find that reducing anthropogenic mortality rates could have a strong beneficial effect on the population: reducing *p_mort_* from the current rate of 0.15 to 0.1 from 2025-2070 resulted in the highest predicted population size in 2070 of 47.0 (Table 1; Fig. 5). However, the extent to which reducing mortality rates to any significant degree may be possible remains in question. Specifically, >72% of caracal mortalities are due to vehicle collisions, the reduction of which may be challenging (Leighton et al., 2022). Likewise, although we find that increasing the rate of natural migration would help minimize the impacts of inbreeding and genetic load (Fig. 5), achieving this by building wildlife corridors through the city of Cape Town appears highly unlikely (Fig. 1, S1). Thus, the remaining and perhaps most feasible option to bolster population viability in the Cape Peninsula is to translocate caracals to the population to initiate a genetic rescue. Genetic rescue has been proven effective in several other instances (Bell et al., 2019; Hedrick & Garcia-Dorado, 2016; Hogg, Forbes, Steele, & Luikart, 2006; Johnson et al., 2010; Whiteley et al., 2015), though remains broadly under-utilized in conservation management (Fitzpatrick et al., 2023; Frankham, 2015; Ralls et al., 2018). Assisted migration may become especially needed in the event that natural migration rates continue to decrease and anthropogenic mortality rates continue to increase (Fig. 5; Table 1). Finally, we also show that genetic rescue is predicted to be effective both when sourcing individuals from the nearby Greater Cape Town population as well as a more divergent population, such as the Central Karoo or Namaqualand population (Fig. 5; Table 1). In fact, we observe somewhat greater beneficial effects when sourcing individuals from a divergent population in terms of reducing levels of inbreeding and genetic load via greater heterosis (Fig. 5; Table 1), due to these populations carrying more divergent haplotypes that are better able to mask recessive deleterious mutations. However, these slight benefits need to be weighed against the possibility of translocating individuals into the Cape Peninsula that are adapted to somewhat different environmental conditions (though see Fitzpatrick et al., 2020), as well as the practical challenges of moving individuals over larger geographic distances.

Several caveats are associated with these simulation results. First, there remains some uncertainty in the genomic deleterious mutation rate parameter, a quantity that appears to have a large influence on model projections (Table 1; Fig. 6). Given that mutation rate estimates do not exist for caracals, we instead used a trio-based estimate for domestic cats (Wang et al., 2022) and assume a coding sequence length derived from the domestic cat genome (Buckley et al., 2020). There are two potential issues with this approach. First, caracals are not domestic cats, thus the validity of using parameters from the domestic cat genome remains uncertain. However, felid genomes appear to be highly conserved (Armstrong et al., 2022; Samaha et al., 2021), suggesting that such parameters may not greatly vary across felids. Another potential issue is that our assumption that deleterious mutations occur only in coding regions ignores the many deleterious mutations that are known to arise in conserved non-coding sequence. However, most of these non-coding deleterious mutations are thought to be weakly deleterious (*s* on the order of 1e-3; (Dukler, Mughal, Ramani, Huang, & Siepel, 2022; Murphy, Elyashiv, Amster, & Sella, 2022; Torgerson et al., 2009)), implying that they likely do not have much of an impact on ecological population dynamics. Another area of uncertainty are dominance parameters, which continue to remain very poorly quantified. Given this uncertainty, we conservatively employ a relatively recessive distribution of dominance coefficients and demonstrate that, if mutations are less recessive, extinction risk may be diminished (Fig. 6). Setting these areas of uncertainty aside, we can derive some assurance that our model is reasonable from the observation that the predicted equilibrium inbreeding load for survival to age 3 of 2*B*=4.67 is broadly consistent with comparable field-based estimates in vertebrates (median 2B=4.5 for survival to sexual maturity; Nietlisbach et al., 2019). This finding illustrates that genomics-based estimates of selection parameters can be reconciled with field-based estimates of inbreeding depression (Kyriazis et al., 2023).

Our results also have implications for caracal populations outside of the Cape Peninsula. Somewhat surprisingly, we find that caracals sampled from the Greater Cape Town, Namaqualand, and Central Karoo populations are moderately inbred, with a mean F_ROH>1Mb_=0.12 that was highly consistent across populations (Figs. 2, S9-S10). This finding was particularly surprising for samples from the Central Karoo and Namaqualand, regions where caracal populations appear to be large and have good connectivity (Avenant et al., 2016). To some extent, this may reflect a legacy of persecution by colonial settlers starting in the 1600s and livestock farmers in the early 20^th^ century (Avenant et al., 2016; van Sittert, 2016). The effects of this colonial legacy on wildlife in South Africa have also been documented in other species, such as the Cape Buffalo (Quinn et al., 2023). However, the abundance of relatively long ROH>1Mb in these populations (Fig. 2, S9-S10), with a mean age of ∼14 generations or ∼42 years (Table S2), suggests that inbreeding has remained ongoing despite growing caracal population densities in these regions over the past several decades (Avenant et al., 2016; Veals et al., 2020). For instance, the individual in our dataset with the highest observed levels of inbreeding (F_ROH_=0.29) and greatest total length of ROH>10Mb (Fig. 2, S9-S10) was sampled from Namaqualand. These continued high levels of inbreeding may be due in part to ongoing persecution of caracals by livestock farmers, who routinely kill caracals in large numbers as a form of predator control (Avenant et al., 2016; Drouilly, Nattrass, & Riain, 2023; Drouilly, Tafani, Nattrass, & O’Riain, 2018; Nattrass, Conradie, Stephens, & Drouilly, 2020). The effect of this persecution, which most greatly impacts naïve younger cats, could be to concentrate caracal reproduction within a select group of adults who have learned to avoid hunting, leading to inbreeding. Thus, these results suggest that even caracal populations that are large and seemingly well-connected may not be immune to the threat of inbreeding.

In conclusion, our work documents the impacts of urban expansion on a wild caracal population, demonstrating that the recent and rapid growth of Cape Town has largely isolated the Cape Peninsula caracal population and resulted in elevated levels of inbreeding. Using simulations, we show that the Cape Peninsula caracal population is predicted to decline in population size over the next ∼50 years, particularly as human pressures continue to mount on the population. Although our results predict a relatively minor risk of extinction in the near future, important exceptions include cases where migration rates continue to decrease, or anthropogenic mortality rates continue to increase (Fig. 5). Ensuring the long-term viability of the Cape Peninsula caracal population is of importance both for the ecological role that predators may play (Avenant et al., 2016; Ripple et al., 2014; Tambling, Avenant, Drouilly, & Melville, 2018) as well as the cultural value of maintaining wild species in the vicinity of a dense urban environment (Güneralp, Lwasa, Masundire, Parnell, & Seto, 2018; Holmes, Rebelo, Dorse, & Wood, 2012). To avert extinction and maintain population viability, we suggest that translocating caracals to initiate a genetic rescue may become necessary in the near future. Our analysis has implications not only for the management of the Cape Peninsula caracal population, but also for the broader discussion of how genomic datasets and computational methods can inform conservation management initiatives such as genetic rescue (Bell et al., 2019; Kyriazis et al., 2023; Shafer et al., 2015; Theissinger et al., 2023). We demonstrate the value of a genomics-informed population viability analyses, employing a state-of-the-art eco-evolutionary simulation tool in SLiM (Haller & Messer, 2019, 2023). Thus, we anticipate this work will help serve as a blueprint for other studies aiming to use computational genomic methods to predict extinction risk in wild populations due to deleterious genetic variation.

## Acknowledgements

C.C.K. and K.E.L. were supported by the National Institutes of Health (R35GM119856 to K.E.L.). We thank the Central Karoo and Namaqualand farmers and professional hunters for allowing access to tissue samples. We thank D. Winterton, J. Broadfield, and numerous volunteers for support during fieldwork. B. Stevens, A. Knight, E. Jordan, and T. Hepburn provided essential veterinarian support during captures. We thank the Cape Leopard Trust, Claude Leon Foundation, the University of Cape Town Research Council, Botanica Wines, Stellenbosch University, the National Research Foundation, Wildlife ACT, the City of Cape Town, Big Cat Rescue, Panthera, and numerous private donors for funding to support L.E.K.S. and fieldwork.

## Data Accessibility

All scripts are available at https://github.com/ckyriazis/caracal_genomics and raw sequence data will be deposited on the Sequence Read Archive upon publication.

## Author Contributions

C.C.K., L.E.K.S, J.M.B., R.K.W., and K.E.L. conceived the study. C.C.K. conducted the analysis and wrote the manuscript with input from all authors. J.M.B., M.D., and S.V. provided samples.

