## Supplemental Information for "The influence of gene flow on population viability in an isolated urban caracal population"

### Supplemental Figures and Tables

**Table S1: Summary of sampled individuals, including population of origin and information on sequencing**

| # | Individual ID | Population | sequencing facility | %mapping | coverage |
|---|---|---|---|---|---|
| 1 | CRTB08 | CK | UCB | 98.83 | 37.06 |
| 2 | CRTB18 | CK | UCB | 98.98 | 25.89 |
| 3 | CRTB20 | CK | UCB | 98.98 | 28.59 |
| 4 | CRTB24 | CK | UCB | 99.12 | 31.65 |
| 5 | MD07 | CK | UCB | 98.74 | 28.96 |
| 6 | MD08 | CK | UCB | 99.01 | 30.87 |
| 7 | MD10 | CK | UCB | 98.7 | 34.36 |
| 8 | MD16 | CK | UCB | 98.8 | 29.29 |
| 9 | MD17 | CK | UCB | 98.5 | 37.07 |
| 10 | MD18 | CK | MedGenome | 98.79 | 38.84 |
| 11 | C23 | CP | MedGenome | 98.07 | 34.37 |
| 12 | CM04 | CP | UCB | 97.95 | 26.17 |
| 13 | CM05 | CP | UCB | 98.94 | 31.91 |
| 14 | CM32 | CP | UCB | 98.04 | 28.047 |
| 15 | TMC02 | CP | MedGenome | 98.86 | 34.63 |
| 16 | TMC06 | CP | UCB | 99.19 | 29.48 |
| 17 | TMC07 | CP | UCB | 99.07 | 29.59 |
| 18 | TMC12 | CP | UCB | 98.74 | 26.76 |
| 19 | TMC16 | CP | UCB | 97.6 | 28.77 |
| 20 | CM09 | GCT | UCB | 99.12 | 29.96 |
| 21 | CM12 | GCT | UCB | 97.71 | 34.45 |
| 22 | CM18 | GCT | UCB | 98.93 | 29.25 |
| 23 | CM29 | GCT | UCB | 99.05 | 28.49 |
| 24 | CM33 | GCT | UCB | 98.81 | 24.99 |
| 25 | TMC20 | GCT | UCB | 98.16 | 27.11 |
| 26 | TMC30 | GCT | UCB | 99.15 | 32.63 |
| 27 | CN01 | NMQ | UCB | 99.16 | 36.94 |
| 28 | CN02 | NMQ | MedGenome | 98.96 | 36.84 |
| 29 | CN03 | NMQ | UCB | 99.03 | 29.28 |
| 30 | CN11 | NMQ | UCB | 99.06 | 43.17 |
| 31 | CN08 | NMQ | UCB | 98.97 | 30.85 |
| | | Average: | | 98.74 | 31.49 |

**Table S2: Estimates of expected age for runs of homozygosity (ROH) for each sampled population**

| Population | Mean ROH>1Mb age (gen) | Mean ROH>1Mb age (years) | Mean ROH>10Mb age (gen) | Mean ROH>10Mb age (years) |
|---|---|---|---|---|
| Cape Peninsula | 10.84 | 32.53 | 2.35 | 7.05 |
| Greater Cape Town | 15.33 | 46.02 | 3.17 | 9.51 |
| Central Karoo | 14.78 | 44.34 | 3.12 | 9.36 |
| Namaqualand | 12.57 | 37.71 | 2.75 | 8.25 |

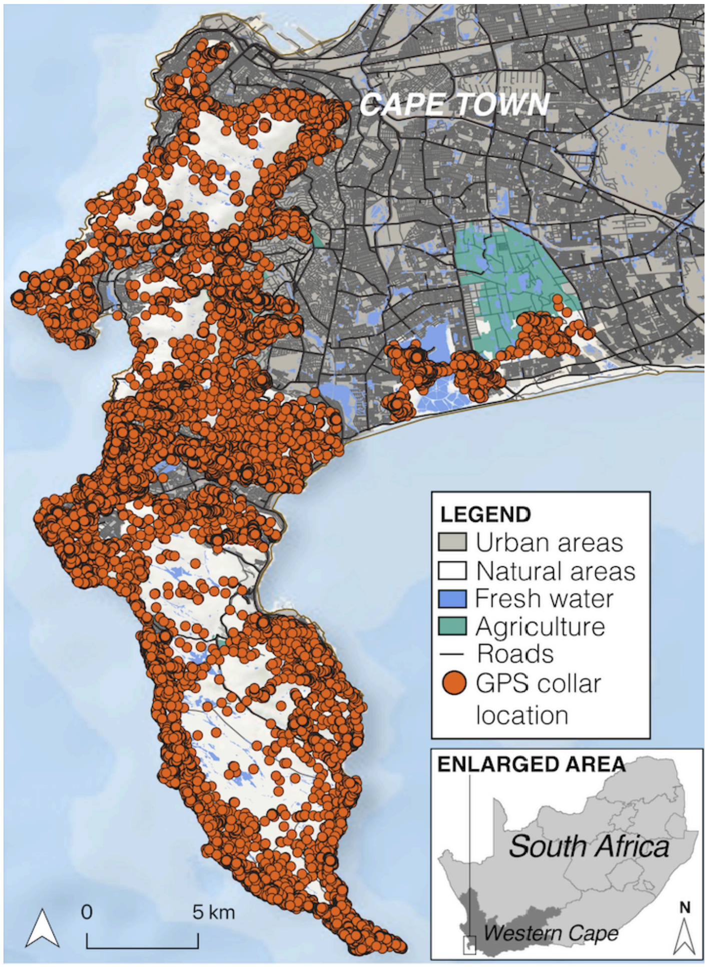


**Figure S1**: **Map of Cape Town including GPS collar data showing how the urban matrix of Cape Town represents a barrier to caracal movement**. Each dot represents the 3-hour GPS collar location for one of 25 caracals. Note that the peninsula is approximately 320 $km^2$ and animals are seen to use nearly all the available green space in the peninsula.

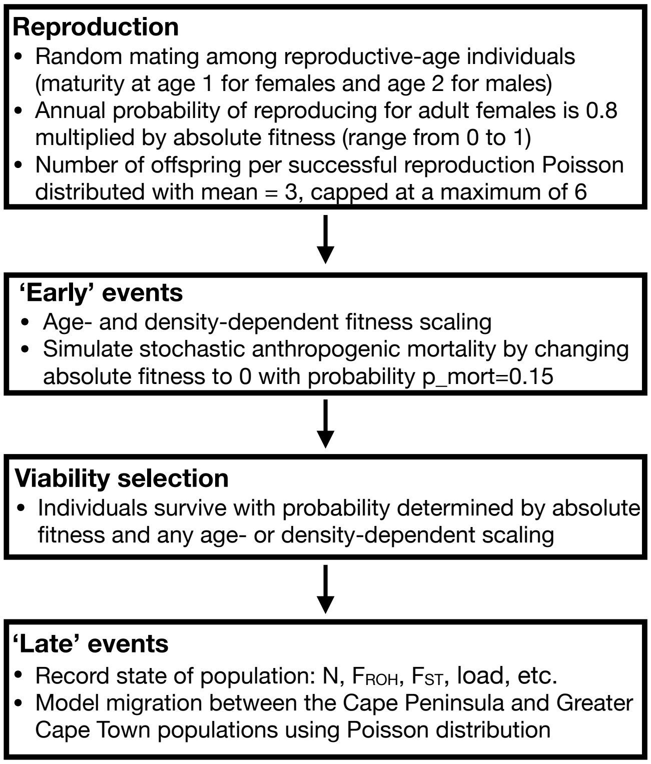


**Figure S2: Schematic of events occurring during one year of the SLiM non-Wright-Fisher simulation model.** See Methods for further details on how the simulation model was parameterized.

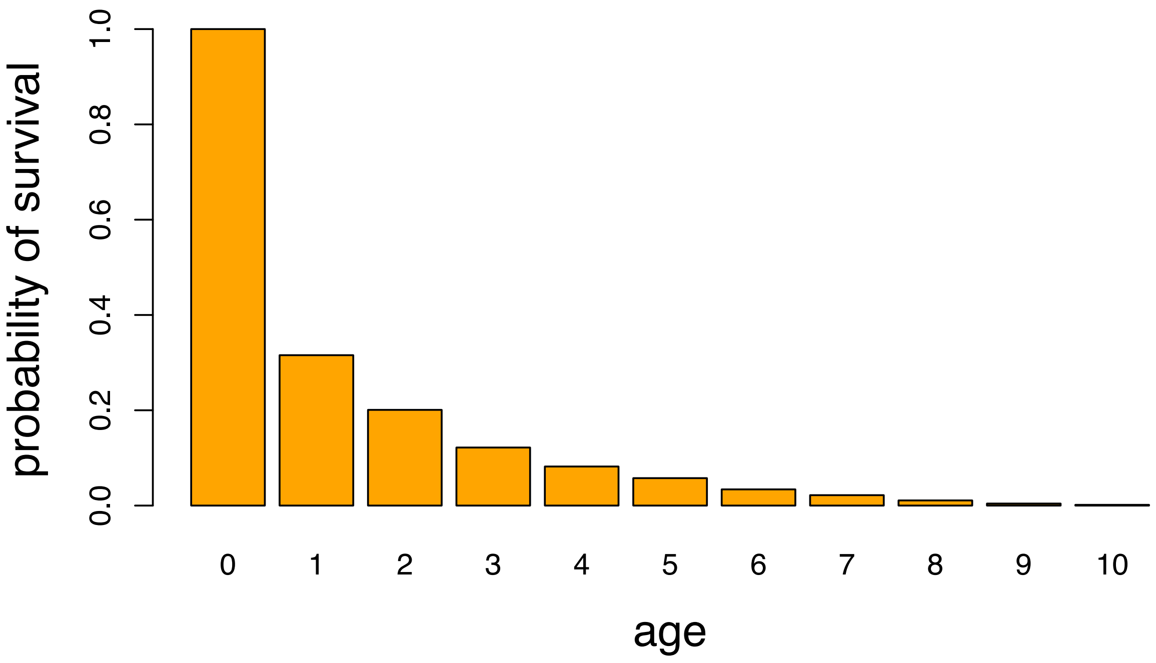


**Figure S3: Survivorship curve for the SLiM simulation model.** Each column represents probability of survival to a given age. Note that ~30% of individuals survive to age 1, and <1% survive to age 10.

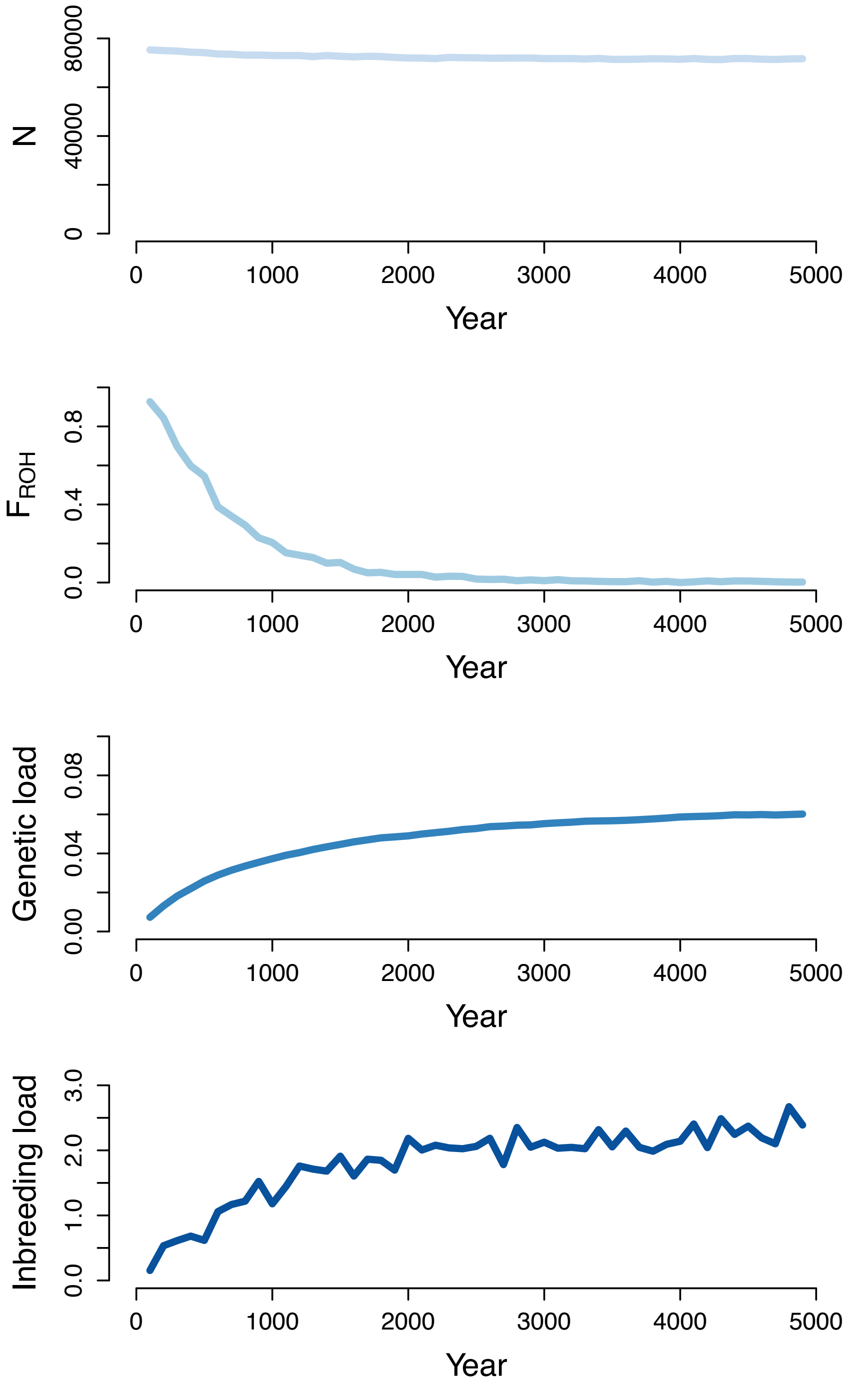


**Figure S4: Burn-in dynamics for the SLiM simulation model.** Plot depicts population size (N), mean inbreeding coefficient ($F_{ROH}$), mean genetic load, and mean inbreeding load (measured as the diploid number of lethal equivalents for annual survival) for the simulated population sampled every 100 years during the burn-in. Note that all quantities approach equilibrium before the end of the burn-in.

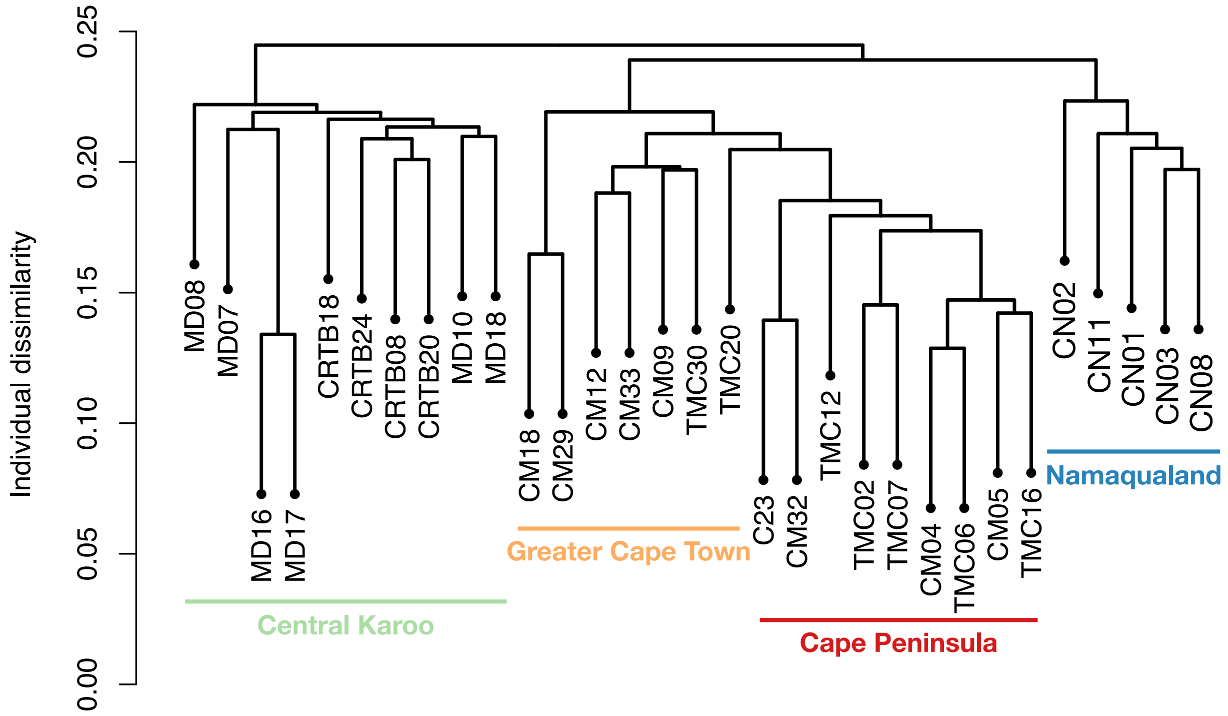


**Figure S5: Hierarchical clustering based on identity-by-state constructed using 49,941 LD-pruned SNPs.** Note that samples from within each population generally cluster together.

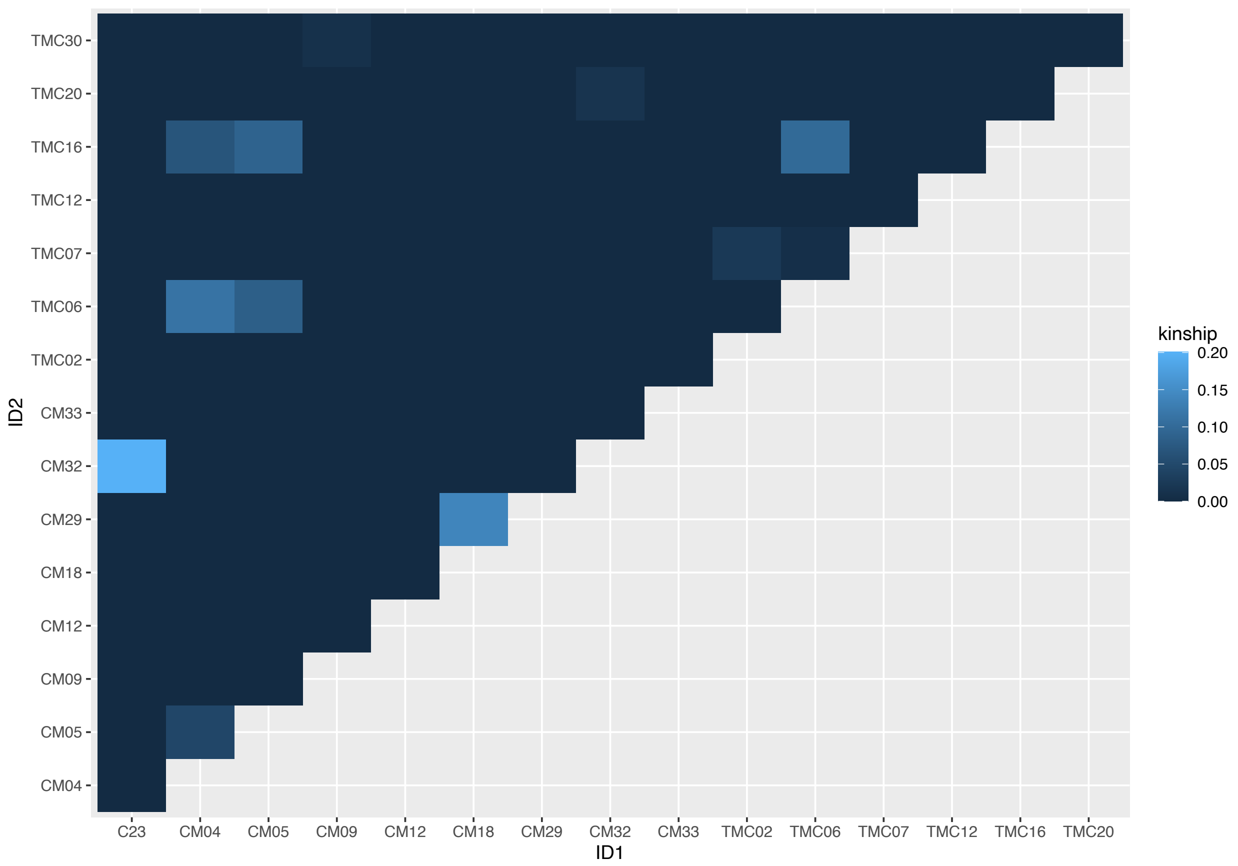


**Figure S6: Heatmap of pairwise kinship estimates for Cape Peninsula and Greater Cape Town individuals.** Note that estimates of kinship coefficients among individuals from other sampled populations were generally close to 0.

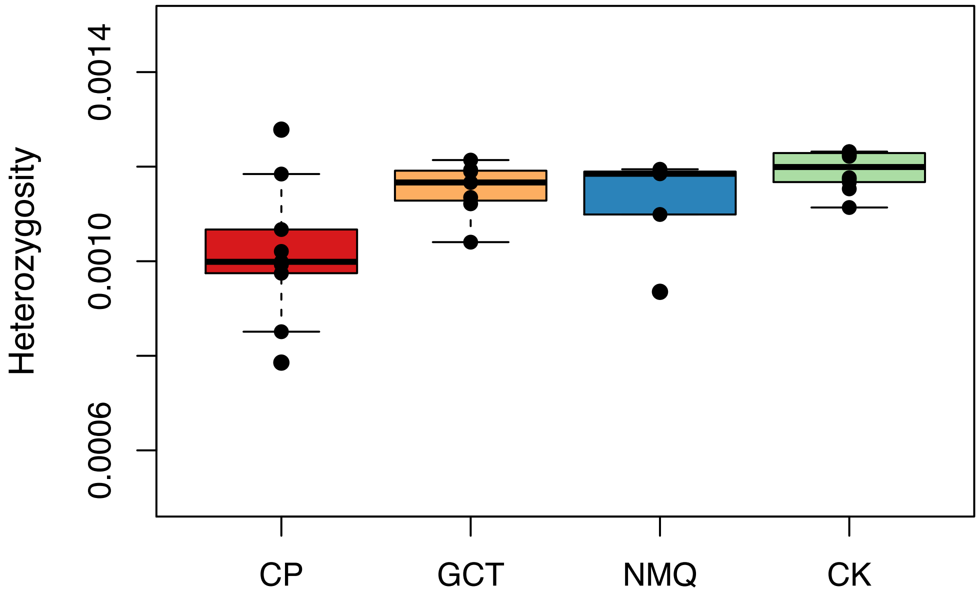


**Figure S7: Boxplots of heterozygosity comparing individuals sampled from four populations.** Note that heterozygosity is much lower in the Cape Peninsula, though is generally consistent across other sampled populations.

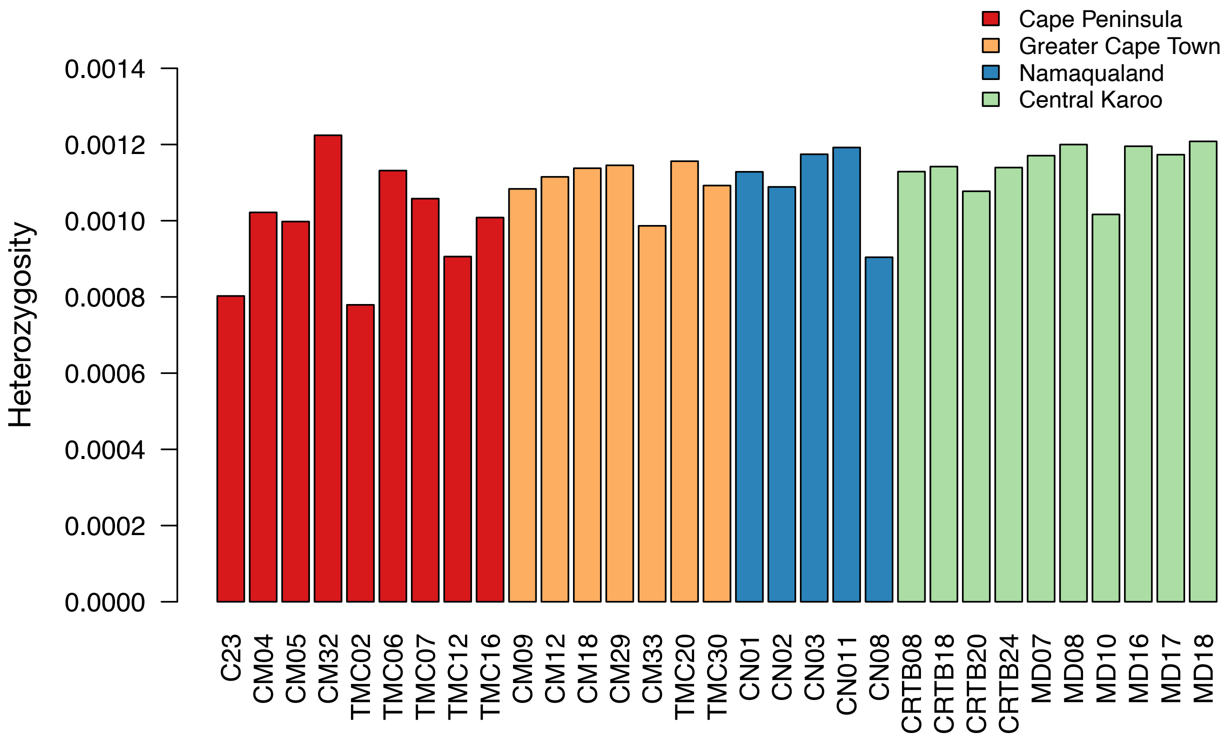


**Figure S8: Heterozygosity estimates for each sampled individual.** Note that heterozygosity is generally lower in the Cape Peninsula, with the main exception of CM32.

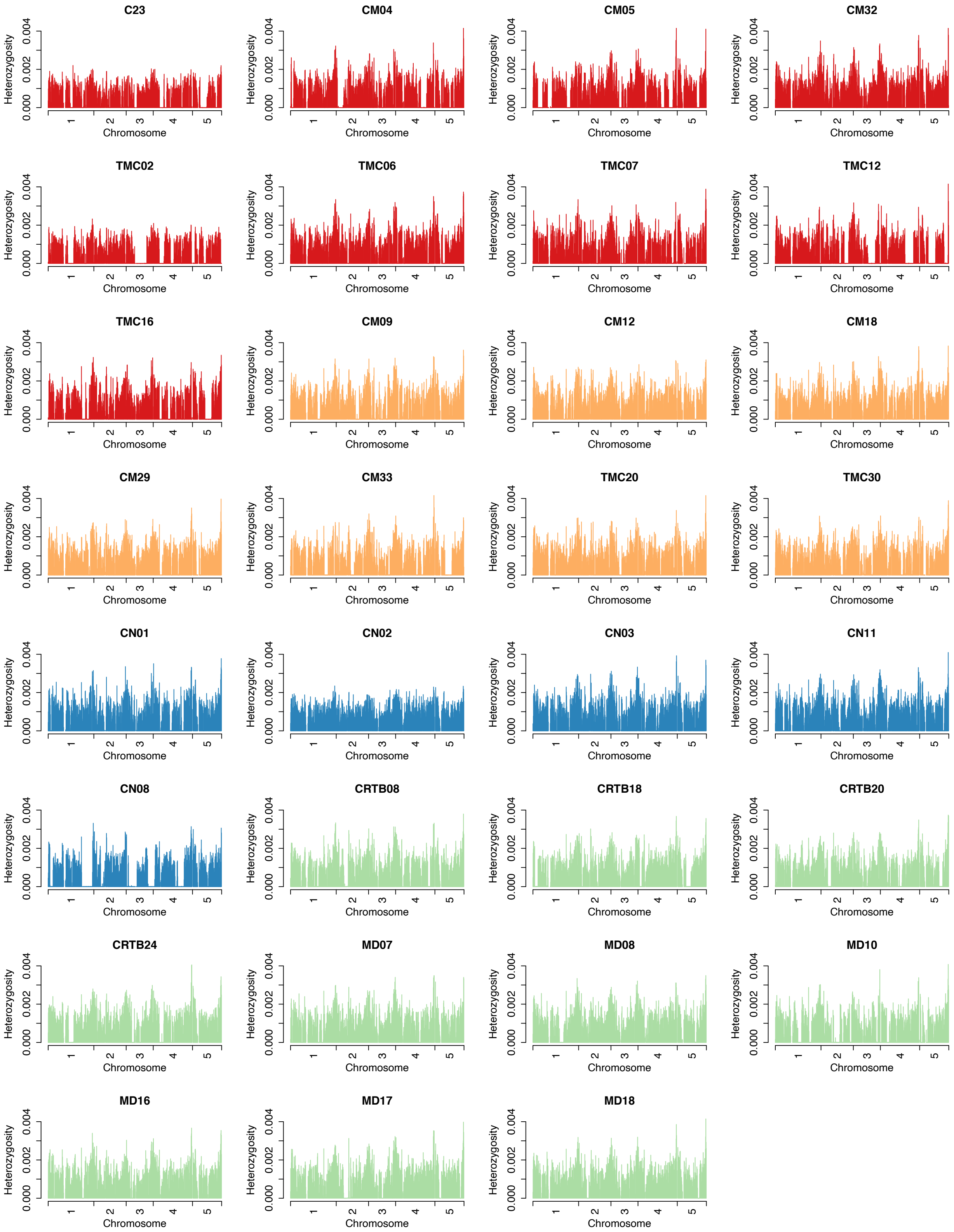


**Figure S9: Plots of heterozygosity in sliding windows for all sampled individuals.** Note the abundance of long ROH in Cape Peninsula individuals, which are largely absent in other sampled individuals. To facilitate visualization, results are shown only for the first 5 chromosomes.

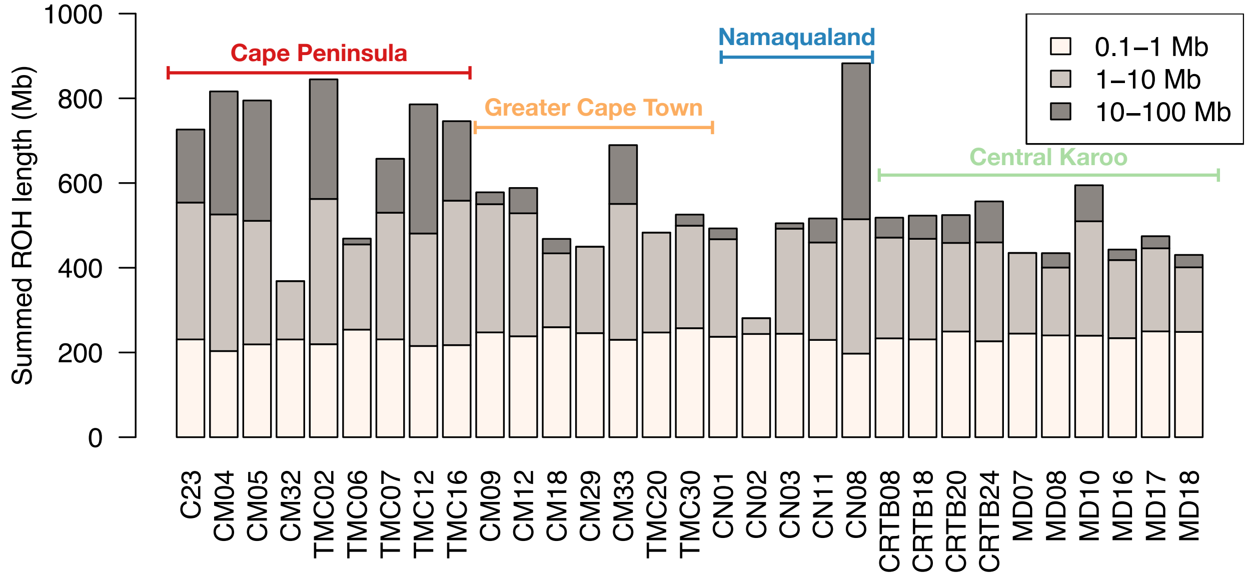


**Figure S10: Summed ROH length for each sampled individual binned into three length categories.** (1) Short ROH 0.1-1 Mb in length, representing shared demographic history. (2) Medium ROH 1-10Mb in length, representing inbreeding over the past ~45 generations or ~135 years. (3) Long ROH >10Mb, representing inbreeding over the past ~5 years or ~15 generations.

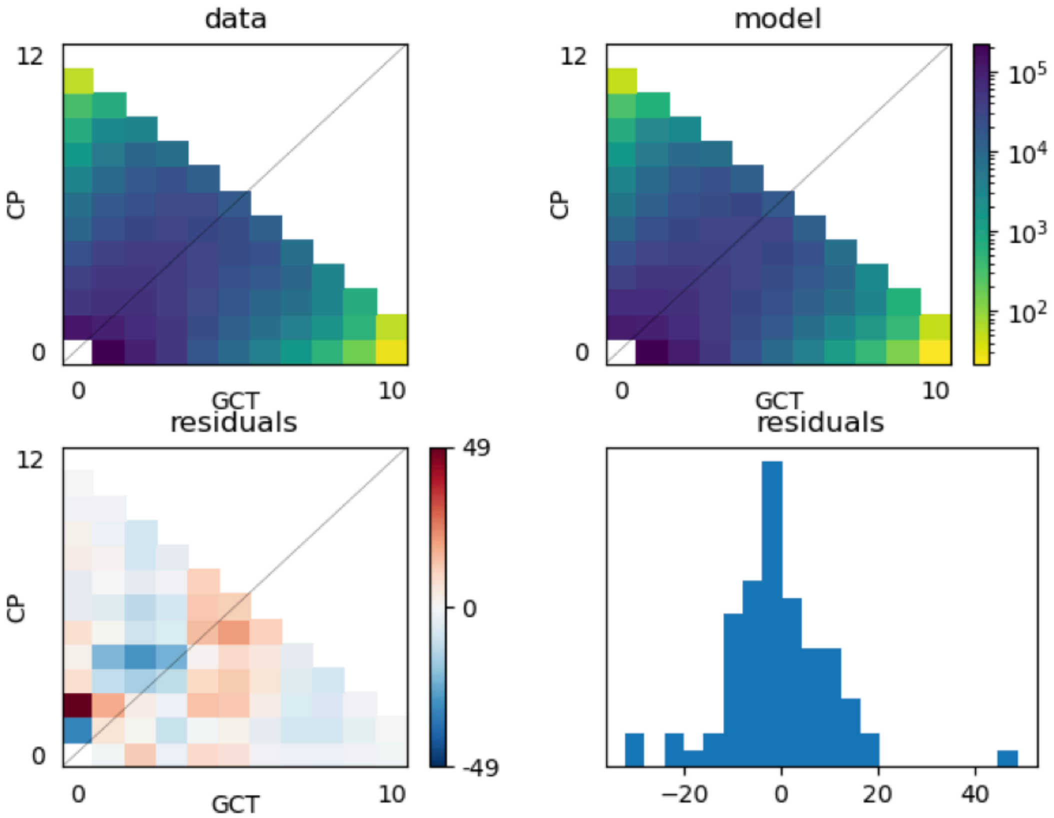


**Figure S11: Plots of the two-dimensional SFS for demographic model inferred using ∂a∂i.** Top panels show 2D SFS observed in data (left) and predicted by model (right) and bottom panels show residuals for each bin (left) and distribution of residuals (right). See Figure 3 for model schematic.

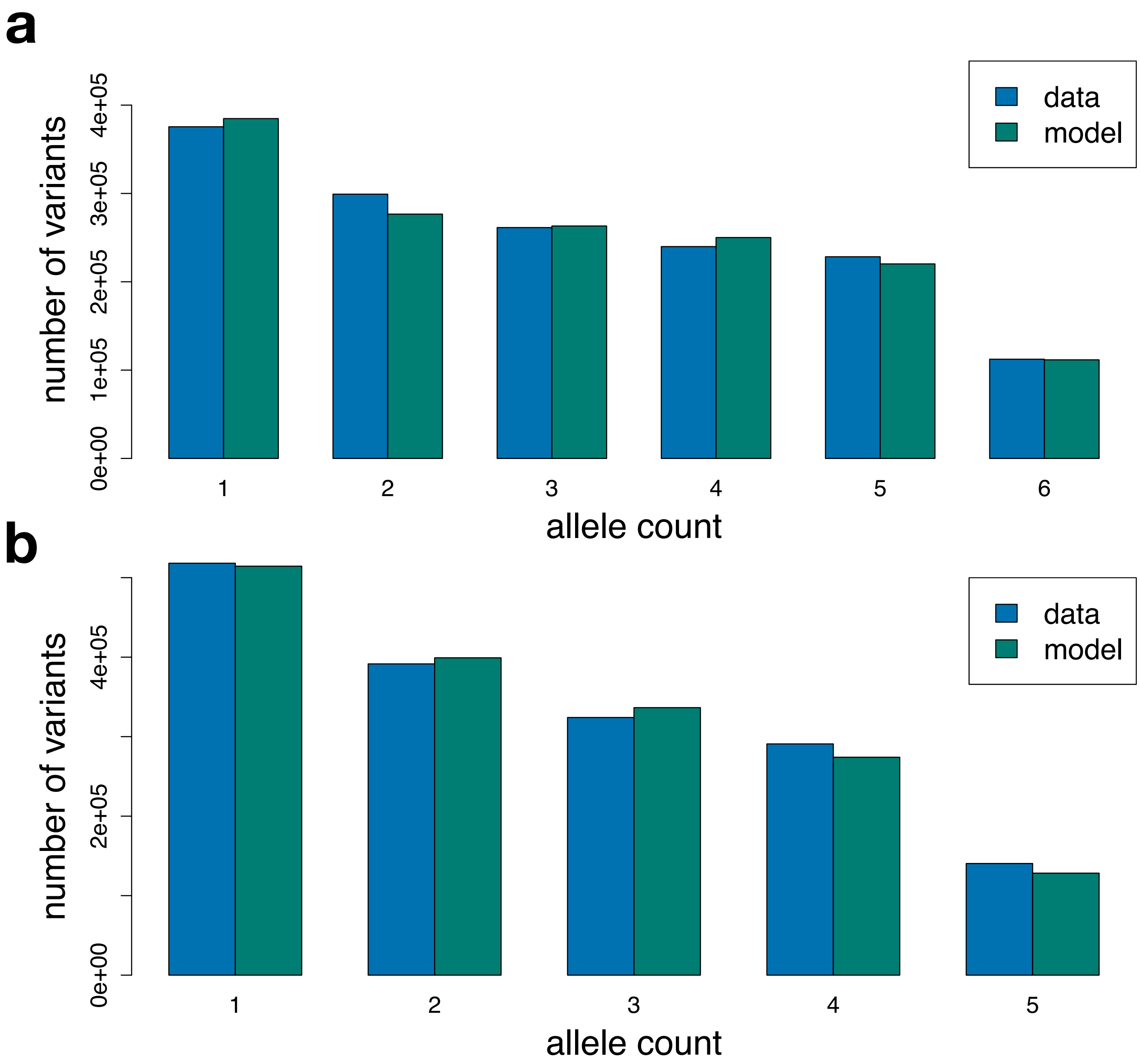


**Figure S12: Plots of marginal one-dimensional SFS for demographic model inferred using ∂a∂i.** (a) Comparison of expected and observed SFS for the Cape Peninsula population. (b) Comparison of expected and observed SFS for the Greater Cape Town population.
